# A single nucleotide polymorphism determines constitutive versus inducible type VI secretion in *Vibrio cholerae*

**DOI:** 10.1101/2022.01.28.478222

**Authors:** Natália C. Drebes Dörr, Alexis Proutière, Milena Jaskólska, Sandrine Stutzmann, Loriane Bader, Melanie Blokesch

## Abstract

*Vibrio cholerae* is a well-studied human pathogen that is also a common inhabitant of marine habitats. In both environments, the bacterium is subject to interbacterial competition. A molecular nanomachine that is often involved in such competitive behavior is the type VI secretion system (T6SS). Interestingly and in contrast to non-pandemic or environmental isolates, the T6SS of the O1 El Tor clade of *V. cholerae*, which is responsible for the ongoing 7^th^ pandemic, is largely silent under standard conditions. Instead, these strains induce their full T6SS capacity only under specific conditions such as growth on chitinous surfaces (signaled through TfoX and QstR) or when the cells encounter low intracellular c-di-GMP levels (TfoY-driven). In this study, we identified a single nucleotide polymorphism (SNP) within an intergenic region of the major T6SS gene cluster of *V. cholerae* that determines the T6SS status of the cell. We show that SNP conversion is sufficient to induce T6SS production in numerous pandemic strains, while the converse approach renders non-pandemic/environmental *V. cholerae* strains T6SS-silent. We further demonstrate that SNP-dependent T6SS production occurs independently of the known T6SS regulators TfoX, QstR, and TfoY. Finally, we identify a putative promoter region adjacent to the identified SNP that is required for all forms of T6SS regulation.

## Main body

Competition between microbes occurs frequently in nature. A widespread molecular nanomachine in Gram-negative bacteria that is involved in interbacterial competition is the type VI secretion system (T6SS). T6SSs resemble inverted contractile phage tails, which, upon contraction, propel a molecular spear out of the bacterium and into neighboring cells. They thereby deliver cocktails of effector proteins for prey intoxication. The producing bacterium and its siblings are protected from self-harm through cognate antitoxins/immunity proteins [1].

The dual lifestyle of *Vibrio cholerae* makes it an ideal model organism to study T6SS-mediated interbacterial competition. Apart from being an important human pathogen, it is also a natural inhabitant of aquatic environments, where it frequently associates with zooplankton and their molted chitinous exoskeletons [2]. Importantly, chitin is a potent inducer of several phenotypes in *V. cholerae* such as natural competence for transformation and type VI secretion [3, 4]. Furthermore, the coupling of competence and T6S fosters the horizontal transfer of prey-released DNA [4, 5].

Chitin-dependent induction of competence and T6SS is triggered through the TfoX signaling pathway, which includes three principal regulatory proteins: TfoX, whose production is tightly linked to growth on chitin [3]; HapR, the master regulator of quorum sensing (QS), which signals high cell density [6]; and the downstream-acting transcription factor QstR [7, 8] (reviewed in [9]). In addition to chitin-dependent regulation, the T6SS is also inducible in *V. cholerae* and other Vibrios by the regulatory protein TfoY [10-12]. However, although TfoY translation occurs at low intracellular c-di-GMP levels [10, 11], the natural trigger for its production remains unknown.

TfoX-, QstR-and TfoY-dependent T6SS induction was primarily demonstrated for *V. cholerae* strains that are responsible for the ongoing 7^th^ cholera pandemic (O1 El Tor strains; referred to as 7PET clade). Indeed, since its discovery, it is well known that the T6SS of 7PET strains is largely silent under laboratory conditions [13]. For this reason, the structural and mechanistic investigation of the T6SS machinery over the past 15 years has primarily relied on constitutively active T6SS strains, using either non-pandemic toxigenic strains (e.g., O37 serogroup strains V52 and ATCC25872) or various environmental isolates [14-16]. In this study, we asked the question “What caused the switch from constitutive to inducible T6SS activity in 7PET strains compared to their non-pandemic or environmental relatives?”.

To address this question, we generated a library of 800 hybrid strains in which each clone carried a mosaic genome between the 7PET pandemic strain A1552 (T6SS OFF) and the non-pandemic strain ATCC25872 (T6SS ON) (Fig. 1A). The library design was based on our previous work on transformation-mediated horizontal gene transfer (HGT) in *V. cholerae* whereby exchanges above 100kb occurred frequently [5]. Specifically, we first created derivatives of strain A1552 bearing an antibiotic resistance marker (*aph*) at 40 different positions throughout its genome, spaced ~100kb apart (Fig. 1A). Isolated genomic DNA of these 40 strains was then used to transform strain ATCC25872, and 20 transformants were isolated per reaction (Fig. 1A). The resulting 800 hybrid transformants were then tested for T6SS activity using a fluorescence imaging-based *E. coli* killing experiment (adapted from [12]). Strikingly, 19/20 hybrid clones that had transferred the *aph* marker at position #32 (*aph*#32) had lost their T6SS activity (T6SS ON>OFF; Fig. S1A). The reverse experiment (i.e., transfer of *aph*#32 from ATCC25872 to A1552) resulted in 7/20 transformants that had gained T6SS activity (T6SS OFF>ON; Fig. S1A). Since the *aph*#32 cassette was located ~15kb upstream of the major/large T6SS cluster (Fig. 1B), we hypothesized that the genomic region that drives constitutive T6SS production might be close to or inside this cluster. Indeed, when we repeated the transfer experiments using strains carrying an insertion (*aph*#42) immediately upstream of *paar1* (first gene in this cluster), this resulted in 20/20 T6SS phenotypic conversion events in the ON>OFF direction and 19/20 events in the OFF>ON direction (Fig. S1A).

**Figure 1:**
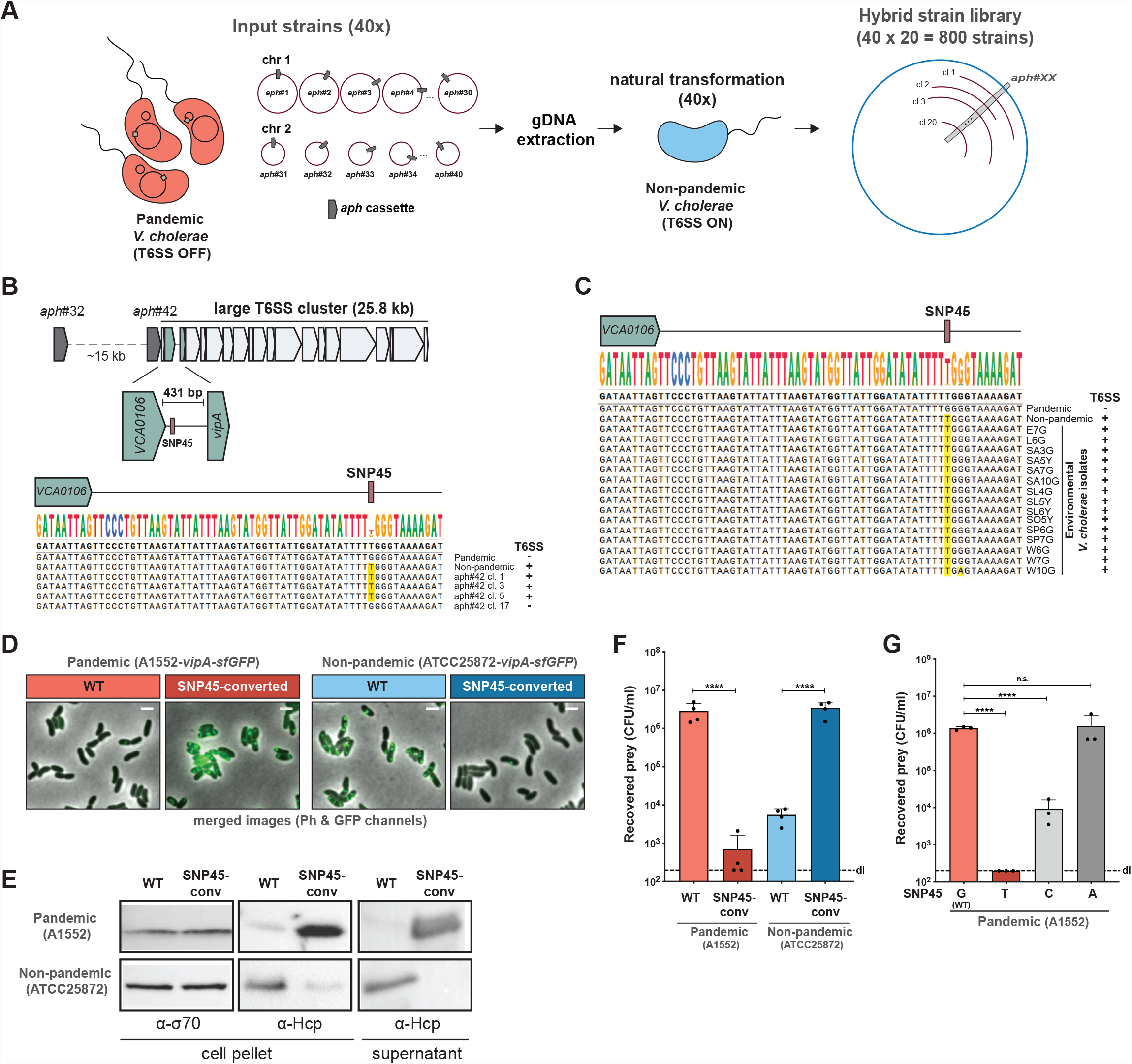
A single SNP determines T6SS activity. **(A)** Scheme of the hybrid strain library construction. 40 input strains in the A1552 strain background (pandemic isolate) were genetically engineered to each carry an *aph* resistance marker at a different genomic location (~every 100kb; *aph*#1 to *aph*#40). Genomic DNA (gDNA) of these strains was used to transform the non-pandemic strain ATCC25872. 20 transformants of each reaction were kept resulting in a final hybrid strain library consisting of 800 strains. **(B)** Scheme of the large T6SS cluster of *V. cholerae* and the location of *aph*#32 and *aph*#42. The zoomed 431-bp intergenic region between the second (*VCA0106*) and third (*vipA*) gene of the cluster is shown below together with an alignment covering the start of this region and comparing the sequence of the pandemic strain A1552, the non-pandemic strain ATCC25872, and four *aph*#42 transformants of strain A1552. The T6SS-ON version of SNP45 (‘T’) is highlighted in yellow and the strains’ T6SS activity status is shown on the right. **(C)** Sequence alignment of the same region as in (B) comparing the pandemic/non-pandemic control strains with 15 environmental *V. cholerae* isolates. **(D-F)** The SNP45-converted pandemic/non-pandemic clones and their parental strains were scored for T6SS assembly (by imaging structures made of the T6SS sheath protein VipA-sfGFP; Scale bars: 2 μm; D), T6SS activity (through Western Blot-based detection of the secreted T6SS tube protein Hcp; E), and interbacterial killing of *E. coli* prey (F). Numbers of surviving prey are depicted on the *Y*-axis (CFU/ml) and the plot represent the average of four independent biological replicates, as indicated by individual dots (±SD). d.l., detection limit. **(G)** SNP45 conversion to T, C, or A in the pandemic strain A1552. Bacterial killing assay as in (F) with three biologically independent replicates. Statistical significance using a one-way ANOVA followed by a Šídák’s multiple comparisons test is indicated comparing each WT with its SNP-converted derivative (F) or each of the SNP45 convertants with the WT (G). ****, *P* < 0.0001; n.s., not significant.

Next, we compared the genome sequences surrounding *aph*#42 in strains A1552 [17] and ATCC25872 (see Supplementary Material and Methods). In addition, we Sanger sequenced the respective region of three T6SS OFF>ON-converted transformants plus the single non-converted T6SS OFF hybrid clone as a negative control. As shown in the alignment (Fig. 1B), this comparison revealed a perfect correlation between the T6SS status and a single nucleotide polymorphism (SNP) at position 45 of the intergenic region, downstream of the second gene of the T6SS cluster (*VCA0106*), whereby ‘G’ resulted in a silenced T6SS, and ‘T’ rendered the transformant T6SS active (Fig. 1B; hereon referred to as SNP45). This finding is strongly supported by the status of SNP45 in 15 environmental *V. cholerae* strains (Fig. 1C) as well as all examined 7PET strains, as was also recently confirmed in a preprint by Ng *et al*. [18].

To prove causality between SNP45 and T6SS activity, we investigated nucleotide 45 using site-directed mutagenesis. SNP45 conversion (G→T) in the pandemic strain A1552 (T6SS OFF) led to expression of the T6SS genes (Fig. S1B and Table S1), assembly of the T6SS machinery, secretion of the T6SS inner tube protein Hcp, and, ultimately, to killing of the *E. coli* prey (Fig. 1D-F). Conversely, SNP45 conversion (T→G) in the non-pandemic strain ATCC25872 (T6SS ON) silenced T6SS activity (Figs. 1D-F, S1C, and Table S2). We confirmed these SNP45 conversion data in five additional 7PET strains (Fig. S2A and B) that were isolated over the past ~40 years from three different continents (Table S3) as well as in a selection of environmental isolates (Fig. S2C and D). Finally, SNP45 conversion (G→C) in strain A1552 resulted in an intermediate activation of T6SS killing, whereas the (G→A) conversion remained T6SS silent (Fig. 1G).

In order to get a first insight into the nature of the SNP45-based regulation, we tested whether the known regulators TfoX, QstR, and TfoY were involved in T6SS induction in the SNP45-converted pandemic strain, which turned out not to be the case (Fig. 2A). We therefore reasoned that an additional regulatory element(s) might be present in the SNP45-harboring intergenic region and therefore stepwise shortened this region (Fig. 2B). Interestingly, while deletion of 276 bp upstream of *vipA* did not impair killing, T6SS activity was completely abolished in a strain lacking -336 bp, despite the presence of SNP45 (Fig. 2C). Consistent with this finding, deletion of the region that differed between these two constructs (60 bp in size) was sufficient to abolish T6SS activity in the SNP45-converted pandemic strain as was the deletion of the almost complete intergenic region (Δfull; Fig. 2C). Visual inspection of the intergenic region surrounding the SNP and encompassing the 60 bp region revealed a putative promoter with appropriately positioned -35 and -10 elements [19] (Fig. 2D). Site-directed mutations designed to disable the -10 element (TAG<u>AA</u>T to TAG<u>GC</u>T) eliminated bacterial killing of the SNP45-converted strain, confirming its importance in T6SS regulation. Importantly, this promoter region was also necessary for TfoX-, QstR-, or TfoY-driven T6SS production in the wild-type pandemic strain A1552 (Fig. 2E).

**Figure 2:**
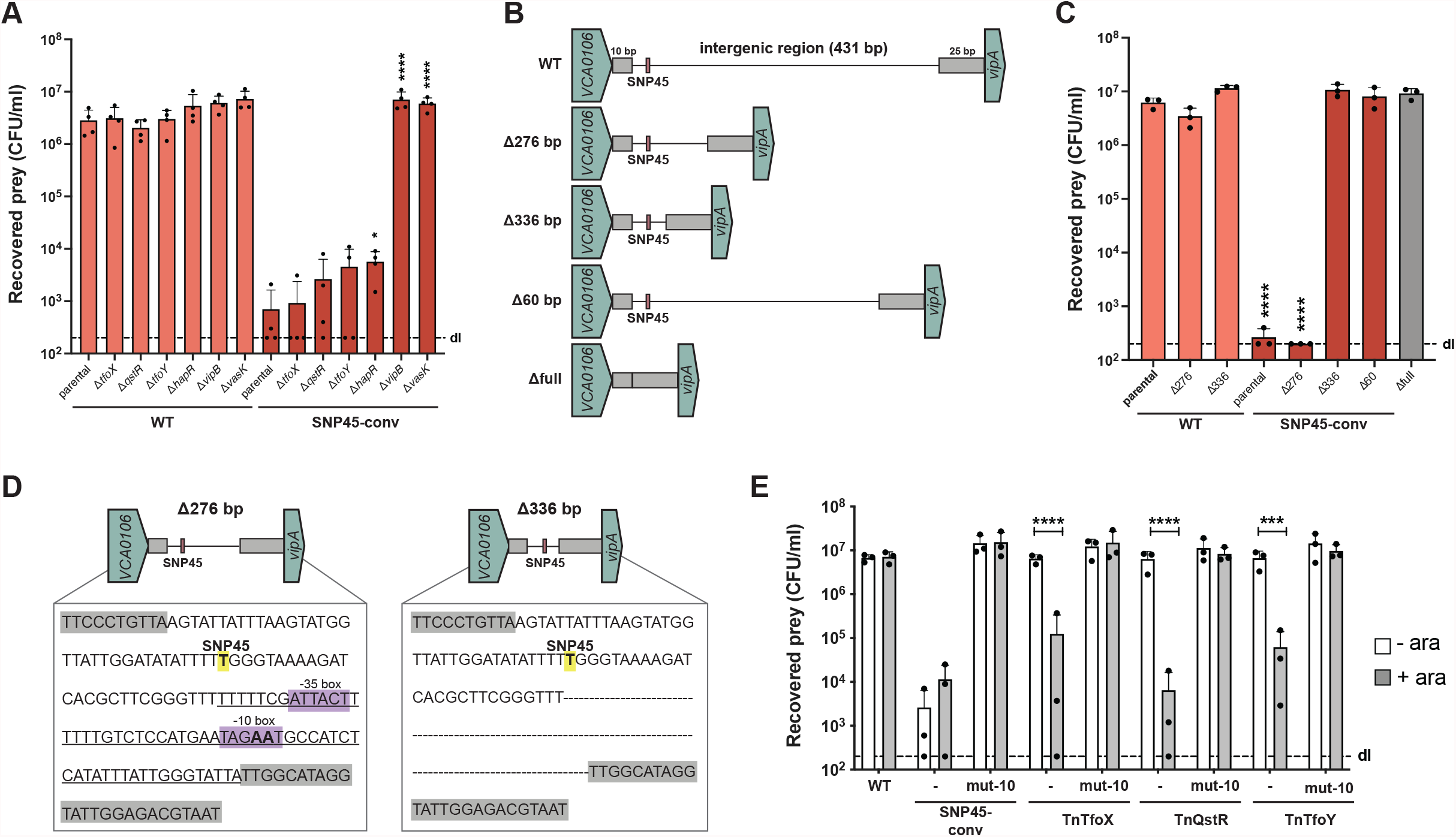
T6SS regulation requires a promoter in the intergenic region. **(A)** Strains lacking known T6SS regulators maintain T6SS activity of the SNP45-converted pandemic strain. Pandemic strain A1552 or its SNP45-converted derivate were genetically engineered to delete *tfoX, qstR, tfoY, hapR*, or the two T6SS structural genes *vipB* and *vasK* as controls. All strains were tested in an *E. coli* killing assay. **(B)** Scheme of truncations introduced within the intergenic region. The intergenic region was shortened by 276 bp, 336 bp or entirely deleted (396 bp deleted; Δfull), leaving solely 10 bp downstream of *VCA0106* and 25 bp upstream of *vipA* intact (gray boxes). **(C)** T6SS activity of strains with a truncated intergenic region as depicted in (B) was assessed in an *E. coli* killing assay. **(D)** A promoter signature is located close to SNP45. Comparison of SNP45-converted (yellow label) intergenic region in the Δ276 and Δ336 mutants with the putative -35 and -10 promoter elements highlighted in purple. The 60bp region deleted in the Δ60 mutant is underlined in the left scheme. The ‘AA’ nucleotides within the -10 element that were changed to ‘GC’ in the respective mutants in (E) are shown in boldface. **(E)** No T6SS activation occurs in strains with a mutated -10 element, as assessed in an *E. coli* killing assay. Neither SNP45-conversion nor arabinose (0.2%)-induced production of TfoX, QstR, and TfoY (from the respective Tn constructs, as described [8, 10]) led to T6SS-mediated prey killing in the mutant strain carrying a defective -10 element (mut-10) in contrast to the WT background. Details for the *E. coli* killing assays in panels (A), (C), and (E) are as described in Fig. 1. Bar plots represent the average of at least three independent biological replicates as shown by the individual dots (±SD). Only statistically significant comparisons are indicated in the plots using one- or two-way ANOVA followed by a Šídák’s multiple comparisons test. The comparisons were: (A) each mutant with its parental strain for WT or SNP45-converted A1552; (C) all strains against the parental WT strain of A1552 shown in boldface; and (E) without or with arabinose conditions for each strain. *, *P* < 0.05; ***, *P* < 0.001; **** *P* < 0.0001.

Collectively, we have identified a SNP in *V. cholerae* that is deterministic of T6SS production. Interestingly, recent work has shown that strains responsible for earlier pandemics, and specifically those belonging to the classical clade of 6^th^ pandemic strains, contain multiple frameshift mutations/deletions in their structural T6SS genes that render them T6SS silent [20]. It is therefore tempting to speculate that the displacement of the classical clade by 7PET clade strains was in part driven by their superior T6SS regulation. Indeed, by keeping their T6SS mostly silent under non-inducing conditions, 7PET strains might keep intestinal inflammation to a minimum [21], while maintaining the ability to produce their T6SS machine “on demand” (e.g., during competition and HGT on chitinous surfaces). Our work will therefore prompt future studies on T6SS regulation in *V. cholerae* and its involvement in a disease context.

## Acknowledgments

The authors thank C. Stoudmann for technical assistance, Eve Rahbé for a transposon mutagenesis screen in strain ATCC25872, A. Vanhove for assisting with the stocking of the hybrid strain library, and members of the Blokesch group for fruitful discussions. The authors also acknowledge the staff of the Lausanne Genomic Technologies Facility at the University of Lausanne for sample processing, PacBio sequencing, and genome assembly and S. Strempel (Microsynth) for the RNA-seq analysis. This work was supported by the Swiss National Science Foundation (310030_185022), the Novartis Foundation for medical-biological Research (#18C178), and a Consolidator grant by the European Research Council (724630). MB is a Howard Hughes Medical Institute (HHMI) International Research Scholar (grant 55008726).

## Author contributions

N.C.D.D. and M.B. conceived the study and designed the experiments; N.C.D.D., A.P., M.J., S.S., L.B., and M.B. performed experiments; M.J. identified the promoter region; N.C.D.D., and M.B. analyzed the data; M.B. secured funding and supervised the study; N.C.D.D. and M.B. drafted the manuscript; A.P. and M.J., commented on the manuscript and all authors approved its final version.

## Additional Information

Supplementary Information. The online version contains supplementary material at https://doi.org/…..

## Content – Supplementary Information

- Supplementary Material and Methods
- Supplementary Figures S1 and S2
- Supplementary Tables S1, S2, S3, S4
- Supplementary References

