## Supplementary Information for "A single nucleotide polymorphism determines constitutive versus inducible type VI secretion in *Vibrio cholerae*"

**Content – Supplementary Information**

- Supplementary Material and Methods
- Supplementary Figures S1 and S2
- Supplementary Tables S1, S2, S3, S4
- Supplementary References

### Supplementary Material and Methods

#### Bacterial strains and growth conditions

Bacterial strains used in this study are listed in Supplementary Table S3. Unless otherwise stated, strains were grown aerobically in Lysogeny broth (LB; 10 g/L of tryptone, 5 g/L of yeast extract, 10 g/L of sodium chloride; Carl Roth) or on LB agar plates at 30°C or 37°C. Half-concentrated defined artificial seawater medium (0.5×DASW) containing HEPES (4-(2-hydroxyethyl)-1-piperazineethanesulfonic acid; Sigma-Aldrich) and vitamins [1] or 0.5× HW Marine Mix (Wiegandt, Germany) were used for growth on chitin (see below for natural transformation-based strain construction). Thiosulfate citrate bile salts sucrose (TCBS, Sigma-Aldrich) agar was used to counter-select *Escherichia coli* following bacterial mating. Counter-selection based on *sacB* was performed on NaCl-free medium containing 10% sucrose. When required, the following antibiotics were added at their given concentrations: kanamycin (75 µg/mL), chloramphenicol (2.5 µg/mL), streptomycin (100 µg/mL), gentamicin (50 µg/mL), and ampicillin (100 µg/mL). To induce expression from the $P_{BAD}$ promoter (inside TnTfoX, TnQstR, and TnTfoY), culture media were supplemented with 0.2% L-arabinose.

#### Recombinant DNA techniques and genetic engineering

DNA manipulations/cloning was carried out using standard methods. PCR amplifications were performed using GoTaq (Promega), Pwo (Roche), or Expand High Fidelity (Roche) polymerases according to the suppliers' recommendations. Genetically modified loci were checked by colony PCR and Sanger sequencing (Microsynth, Switzerland).

*V. cholerae* were genetically engineered using a set of different methods. Natural transformation on chitin flakes and FLP-recombination (TransFLP; [2, 3]) were used to integrate the *vipA-sfGFP* translational fusion construct allele into the chromosome of diverse strains as reported [4]. Natural transformation on chitin flakes using PCR fragments was also used for the hybrid strain library construction (see details below). All remaining mutations and deletions were done by allelic exchange using the counter-selectable plasmids pGP704-Sac28 and pGP704-Sac-Kan [5, 6].

**Construction of a pandemic/non-pandemic hybrid strain library**

The kanamycin-resistance marker *aph* was integrated by natural transformation into the genome of pandemic *V. cholerae* strain A1552. To do so, amplified PCR fragments served as transforming material, whereby each fragment combined an upstream region, *aph* preceded by its promoter, and a downstream region (fused using overlapping PCR). The PCR fragments were prepared to generate 40 transformants (*aph*#1 to *aph*#40) whereby each strain contains the *aph* cassette at a different location (roughly every 100 kb). Genomic DNA (gDNA) of these 40 donor strains was isolated from 2 ml of an overnight culture using 100/G Genomic-tips together with a Genomic DNA buffer set (Qiagen) as described in the manufacturer's instructions.

To generate the pandemic/non-pandemic hybrid strain library, the acceptor strain ATCC25872 was grown in 40 parallel tubes on chitin flakes and each donor strain-derived gDNA was added to one of those tubes. After incubation at 30°C for 30 h, bacteria were selected on LB plates containing kanamycin and 20 transformants were isolated and stocked from each independent reaction, resulting in 800 strains in total.

After the first screen of the 800 strains library, the reverse experiment from what was described above was performed, generating a set of ATCC25872 strains carrying *aph* at 40 different genomic locations. However, transformation of A1552 as acceptor strain was only performed using gDNA of strain ATCC25872-*aph*#32 as transforming material. 20 transformants of this reaction were isolated and stocked and further explored for their T6SS activity. Based on the screening result and the proximity of *aph*#32 to the large T6SS cluster, an additional construct was generated in both strains (A1552-*aph*#42 and ATCC25872-*aph*#42) and their gDNA used to transform in each case the opposite strain to obtain 2 x 20 additional hybrid clones.

**Phenotypic screening of the hybrid strain library**

The initial library was screened for the strains' T6SS activities using a fluorescence-based *E. coli* killing experiment, which was adapted from [7]. Briefly, 200 µL of the predator hybrid strain were mixed with 40 µL of GFP-labelled *E. coli* prey (strain MC4100-TnGFP) in 96-well plates and 5 µL of the mixtures were spotted onto LB plates. The donor (A1552; T6SS OFF) and acceptor (ATCC25872; T6SS ON) strains served as controls. After 4 h incubation at 37°C, the plates were observed under a stereo microscope (Leica EL6000) equipped with a green fluorescence (FITC) filter cube. Prey survival was scored based on their maintained GFP signal. To properly quantify T6SS activity, all *aph*#32 and *aph*#42 transformants were rechecked using the standard interbacterial killing assay described below with *E. coli* TOP10-TnGFP ($Cm^R$) as prey.

**Interbacterial killing assay**

Bacterial killing was assessed following a previously established assay [4]. Briefly, the respective predator and the *E. coli* prey were mixed at a ratio of 10:1 and spotted onto filters on prewarmed LB agar plates (containing 0.2% arabinose where indicated in the figure legend). After 4 h of incubation at 37°C, the bacteria were resuspended, serially diluted, and spotted onto antibiotic-containing (streptomycin, kanamycin, or chloramphenicol) LB agar plates to enumerate colony-forming units (shown as log-transformed CFU/mL in the graphs). Experiments were performed at least three independent times. Statistically significance was determined using using GraphPad Prism 9.1.1 (for macOS) on log-transformed data [8] using a one- or two-way ANOVA followed by a Šídák's multiple comparisons test, as indicated in the figure legends. If no prey bacteria were recovered, the detection limit was used to calculate the mean of the independent experiments and to perform the statistical analysis.

**Imaging of T6SS sheath structure**

To image T6SS sheath structures in strains carrying the translational fusion *vipA-sfGFP*, the bacteria were mounted on microscope slides coated with a thin agarose pad (1.2% in 0.5X PBS), covered with a coverslip, and observed in the phase contrast and epifluorescence mode (green channel) using a Zeiss LSM 700 inverted confocal laser scanning microscope with an attached HXP 120 light unit (Zeiss, Feldbach, Switzerland). Images were adjusted for contrast and brightness and cropped using the Fiji software [9]. The images are overlays of the Ph and GFP channels and representative of at least three biologically independent replicates.

**Whole-genome sequencing and *de novo* assembly of strain ATCC25872**

Bacterial growth and gDNA extraction were done as described [10]. DNA sample preparation and genome sequencing was performed by the Genomic Technology Facility of the University of Lausanne (Switzerland). Briefly, the DNA sample was sheared in Covaris g-TUBEs resulting in fragments with a mean length of 20 kb. The DNA library was prepared using the PacBio SMRTbell template prep kit 1 (Pacific Biosciences) according to the manufacturer's recommendations and size selected on a BluePippin system (Sage Science, Inc.) for molecules larger than 15 kb. The library was sequenced on a PacBio system within one single-molecule real-time (SMRT) cell with P6/C4 chemistry and MagBeads at a movie length of 360 min. The genome was *de novo* assembled using the protocol RS_HGAP_Assembly.3 in SMRT Pipe 2.3.0 and circularized using the Minimus assembler of the AMOS software package 3.1.0 using default parameters [11]. The assembled genome was annotated using Prokka 1.12 [12] but due to incompatibly with the NCBI database, reannotated using their Prokaryotic Genome Annotation Pipeline (PGAP) during data submission. The data are available under BioSample SAMN13736322 and BioProject PRJNA599000 and the NCBI accession numbers CP047305 (chromosome 1) and CP047306 (chromosome 2). Details on the genome sequencing and assembly are provided in Supplementary Table S4.

**Expression profiling by RNA sequencing**

Overnight cultures of strains A1552 and ATCC25872 and their SNP45-converted derivatives were back-diluted 1:100 in LB medium and grown for 3 h at 30°C with agitation. Cells were harvested by centrifugation at 4°C and washed with PBS buffer, followed by lysis with Tri Reagent (Sigma-Aldrich) and shock freezing in a dry-ice ethanol bath. The samples were stored at -80 °C prior to processing. RNA preparation and DNase treatment were performed as previously described [13]. After DNase treatment, an additional purification step was performed using the GenElute Mammalian Total RNA miniprep kit (Sigma-Aldrich).

Downstream processing of the samples was performed by Microsynth AG (Balgach, Switzerland) who also analyzed the data. Briefly, Illumina's TruSeq Stranded Total RNA Library Prep Gold kit including ribodepletion was used to construct libraries from total RNA. Subsequently, an Illumina NextSeq 550 platform and a high output v2.5 kit were used to sequence the library. Single end reads (1x75bp), which passed Illumina's chastity filter, were demultiplexed and trimmed off the Illumina adaptors using Illumina's bcl2fastq software version v2.20.0.422 (no further refinement or selection). The quality of the reads in fastq format was checked with the FastQC software (version 0.11.8) (http://www.bioinformatics.babraham.ac.uk/projects/fastqc/). The reads were mapped to the reference genomes (A1552 chromosome 1 [CP028894] and 2 [CP028895]; ATCC25872 chromosome 1 [CP047305] and 2 [CP047306]) via bowtie2 (version 2.3.5.1) [14] in local mapping mode with very sensitive pre-settings. To count the uniquely mapped reads to annotated genes, the software htseq-count (HTSeq version 0.11.2) [15] was used. Normalization of the raw counts and differential gene expression analysis was carried out with help of the R software package DESeq2 (version 1.22.2) [16]. Data were visualized using the Integrative Genomics Viewer [17].

**Quantitative Reverse Transcription PCR (qRT-PCR)**

To analyze gene expression using quantitative reverse transcription PCR (qRT-PCR), overnight cultures were back-diluted 1:100 in LB medium and grown for 3 h at 30°C with agitation. RNA purification, DNase treatment, cDNA synthesis and qPCR followed a previously established protocol [13]. Samples were analyzed on a LightCycler Real-Time PCR System (LightCycler Nano or LightCycler 96; Roche) using the standard curve method. Expression values are presented relative to the transcript levels of the reference gene *gyrA*. Each experiment was performed two independent times.

**Western blotting**

To check the production of the inner tube T6SS protein Hcp, cell lysates were prepared as described previously [18]. In brief, overnight cultures were back-diluted 1:100 in LB medium and grown with agitation at 30°C for 3 h. Cells were harvested by centrifugation and the bacterial pellet was resuspended in 2× Laemmli buffer (Sigma-Aldrich), adjusting for the total number of bacteria according to the cultures' optical density at 600nm ($OD_{600}$) values. To check for T6SS-secreted Hcp, 1.5 ml of the culture supernatant was filter sterilized (0.2 µm filter; VWR) and the proteins were precipitated using trichloroacetic acid (TCA). The precipitated proteins were washed with acetone before resuspension in 30 µl of 2× Laemmli buffer (Sigma-Aldrich). All samples were heated at 95°C for 15 min.

Proteins were separated by sodium dodecyl sulfate (SDS) polyacrylamide gel electrophoresis (PAGE) using 15% gels and then blotted as previously described [13]. Primary antibodies against Hcp (raised against synthetic peptides, Eurogentec #1510528; [18]) were diluted 1:5,000 while the Anti-rabbit IgG HRP (Sigma-Aldrich, A9169) secondary antibody was diluted 1:20,000. Loading controls were performed using the anti-Sigma70-horseradish peroxidase (HRP) conjugate (BioLegend, 663205) at a 1:10,000 dilution. Lumi-Light$^{PLUS}$ Western Blotting Substrate (Roche, Switzerland) served as the HRP substrate. The signal was detected using a ChemiDoc XRS+ station (BioRad).

### Supplementary Figures

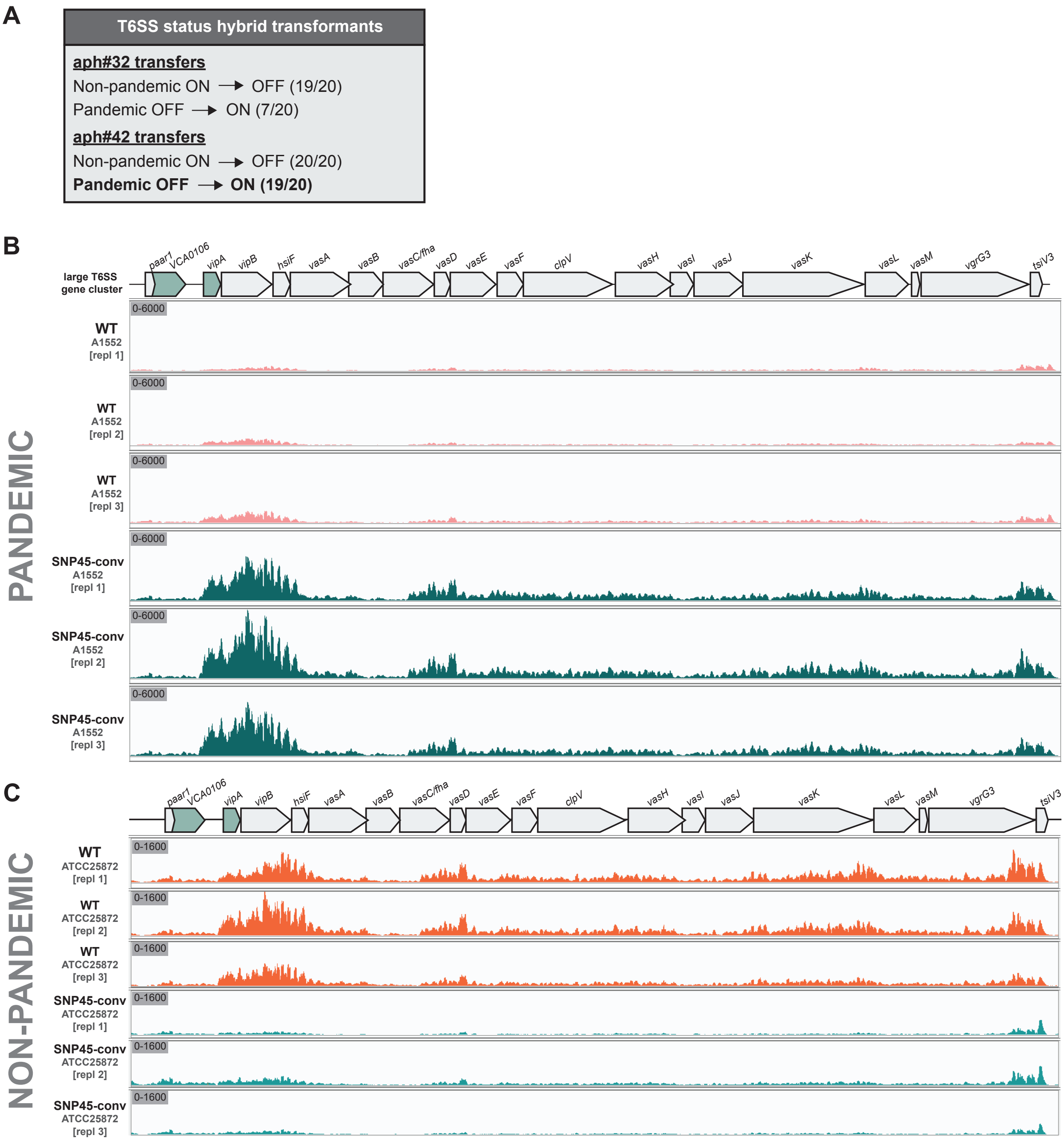


**Figure S1. SNP45-conversion changes T6SS-related phenotypes. (A)** Hybrid strains with changed T6SS activity status. Table summarizes the number of *aph*#32 and *aph*#42 transformants that changed their T6SS phenotype compared to the parental strain (non-pandemic strain ATCC25872 or pandemic strain A1552). The single non-converted transformed clone #17 in Fig. 1B was part of the set highlighted in bold. **(B-C)** SNP-conversion leads to changed transcript abundance. RNA-sequencing data with normalized reads matched to the large T6SS cluster region (*paar1* to *tsiV3*) of *V. cholerae* A1552 (B) or ATCC25872 (C). Three biologically independent replicates of WT and their SNP45-converted derivatives are shown. The scales of the *Y*-axis are shown on the upper left of each row. See Supplementary Tables S1 and S2 for details.

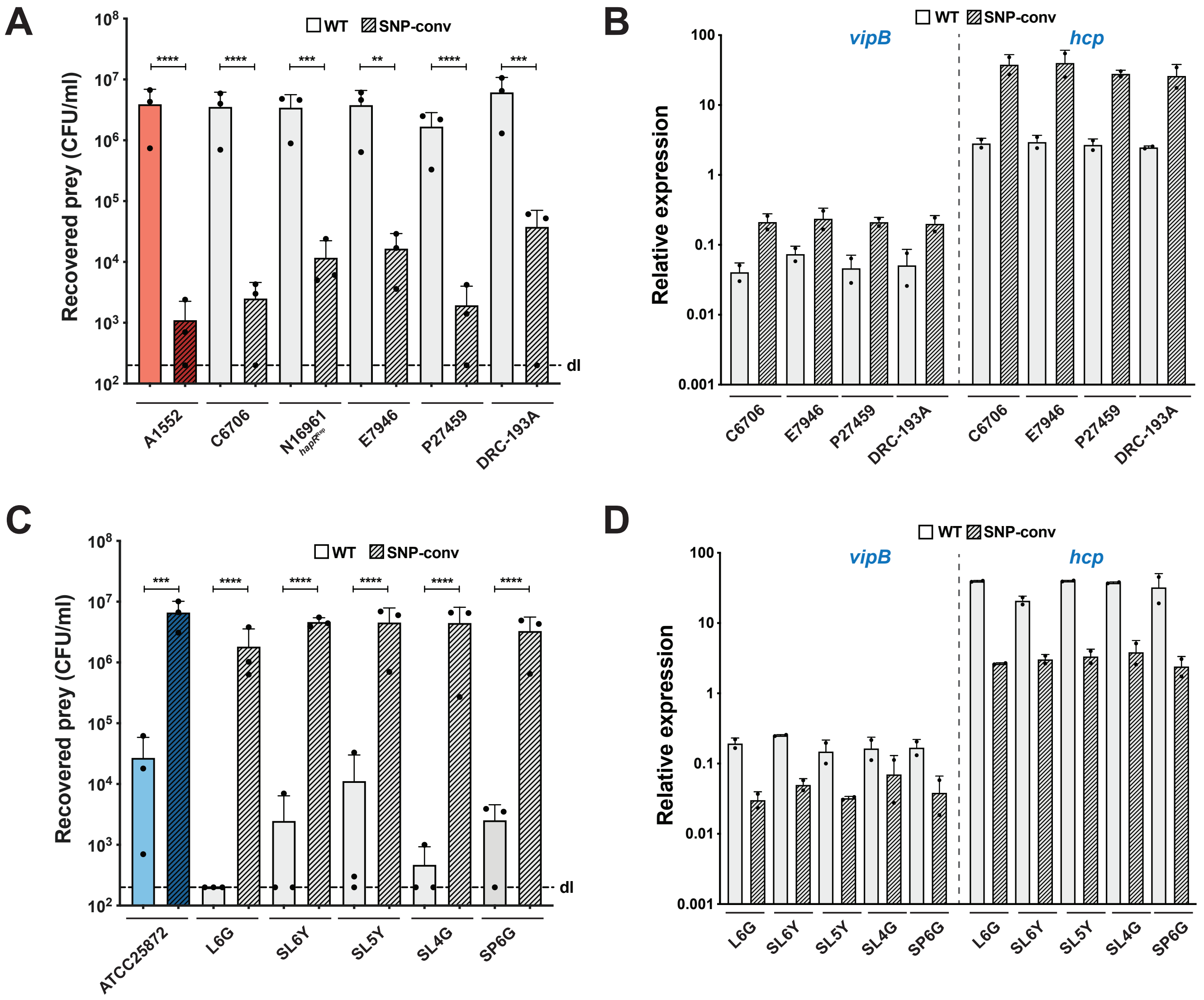


**Figure S2. SNP45-conversion changes T6SS status in pandemic and environmental strains.** SNP45-conversion inverts the ability of strains to kill prey bacteria, as assessed in an *E. coli* killing assay. Results for WT and SNP45-converted derivatives of six well-studied pandemic strains (N16961 derivative is QS-repaired; [24]) **(A)** or five environmental isolates (with ATCC25872 as control) **(C)** are shown. d.l., detection limit. Bar plots represent the average of three independent biological replicates as shown by the individual dots (±SD). Statistically analysis was done using one-way ANOVA followed by a Šídák's multiple comparisons test whereby WT and SNP-converted derivatives were compared. **, $P < 0.01$; ***, $P < 0.001$; ****, $P < 0.0001$. **(B, D)** A selection of the strains from (A, C) were scored for the expression of two conserved T6SS genes (*vipB* and *hcp*) using qRT-PCR. Data represent the average of two independent biological replicates (±SD). Details are provided in the Supplementary Methods section.

**Table S1.** RNA-seq expression data for the T6SS gene cluster in WT and SNP45-converted strain A1552.

| cluster names / comments | gene names | ID (locus tag) in strain A1552[*] | Homologous locus tag in strain N16961[#] | A1552 (Exp1) | A1552 (Exp2) | A1552 (Exp1) | AVG - A1552 | A1552-SNP45 converted (Exp1) | A1552-SNP45 converted (Exp2) | A1552-SNP45 converted (Exp3) | AVG - A1552-SNP45 converted | Fold induction [A1552-SNP45 converted / A1552] |
|---|---|---|---|---|---|---|---|---|---|---|---|---|
| major T6SS regulators | ***tfoX*** | A1552VC_00900 | VC1153 | 7026.857 | 4415.423 | 5957.340 | 5799.87 | 4017.74 | 4330.16 | 5871.61 | 4739.83 | **0.82** |
| | ***qstR*** | A1552VC_00153 | VC0396 | 148.984 | 237.743 | 196.751 | 194.49 | 148.98 | 237.74 | 196.75 | 194.49 | **1.00** |
| | ***tfoY*** | A1552VC_01512 | VC1722 | 8108.02 | 5598.86 | 5787.62 | 6498.17 | 6959.09 | 5845.20 | 5733.44 | 6179.24 | **0.95** |
| Auxiliary cluster 1 | ***hcp1*** | A1552VC_01148 | VC1415 | 3888.62 | 3752.38 | 4343.53 | 3994.85 | 68835.24 | 81594.59 | 83017.62 | 77815.82 | **19.48** |
| | ***vgrG1*** | A1552VC_01149 | VC1416 | 1037.42 | 1096.26 | 1296.90 | 1143.53 | 7124.82 | 14324.05 | 9930.36 | 10459.74 | **9.15** |
| | ***tap1*** | A1552VC_01150 | VC1417 | 807.79 | 763.42 | 831.31 | 800.84 | 4517.61 | 7660.83 | 5163.16 | 5780.54 | **7.22** |
| | ***tseL*** | A1552VC_01151 | VC1418 | 3377.43 | 2611.21 | 2666.65 | 2885.10 | 14045.59 | 22291.06 | 15230.76 | 17189.14 | **5.96** |
| | ***tsiV1*** | A1552VC_01152 | VC1419 | 717.58 | 515.11 | 563.97 | 598.89 | 2691.86 | 4324.57 | 3247.37 | 3421.27 | **5.71** |
| Auxiliary cluster 2 | ***hcp2*** | A1552VC_A02810 | VCA0017 | 4202.99 | 4275.42 | 5147.06 | 4541.82 | 75355.93 | 92646.59 | 97758.78 | 88587.10 | **19.50** |
| | ***vgrG2*** | A1552VC_A02811 | VCA0018 | 225.53 | 339.44 | 357.46 | 307.48 | 2694.53 | 4129.15 | 3360.50 | 3394.73 | **11.04** |
| | ***vasW*** | A1552VC_A02812 | VCA0019 | 299.34 | 320.95 | 340.18 | 320.16 | 2709.68 | 3929.06 | 3029.46 | 3222.73 | **10.07** |
| | ***vasX*** | A1552VC_A02813 | VCA0020 | 2476.69 | 2119.88 | 2134.97 | 2243.85 | 16716.95 | 23313.80 | 17474.81 | 19168.52 | **8.54** |
| | ***tsiV2*** | A1552VC_A02814 | VCA0021 | 1003.25 | 648.51 | 737.44 | 796.40 | 4875.81 | 4819.66 | 4605.86 | 4767.11 | **5.99** |
| Large/major cluster | ***paar1*** | A1552VC_A02890 | VCA0105 | 373.14 | 311.71 | 344.69 | 343.18 | 632.64 | 587.22 | 656.52 | 625.46 | **1.82** |
| | ***VCA0106*** | A1552VC_A02891 | VCA0106 | 512.56 | 624.74 | 687.88 | 608.39 | 1523.69 | 1676.04 | 2070.64 | 1756.79 | **2.89** |
| | ***vipA*** | A1552VC_A02892 | VCA0107 | 893.90 | 2135.73 | 2047.11 | 1692.25 | 7707.57 | 11145.99 | 10666.63 | 9840.06 | **5.81** |
| | ***vipB*** | A1552VC_A02893 | VCA0108 | 4528.30 | 8973.49 | 8484.31 | 7328.70 | 36769.10 | 53117.73 | 44061.18 | 44649.34 | **6.09** |
| | ***hsiF*** | A1552VC_A02894 | VCA0109 | 1097.56 | 1392.12 | 1420.06 | 1303.25 | 7324.41 | 8946.02 | 7155.91 | 7808.78 | **5.99** |
| | ***vasA*** | A1552VC_A02895 | VCA0110 | 1264.31 | 1701.19 | 1619.81 | 1528.44 | 7924.09 | 10588.55 | 8130.49 | 8881.05 | **5.81** |
| | ***vasB*** | A1552VC_A02896 | VCA0111 | 371.78 | 392.28 | 487.37 | 417.14 | 1976.34 | 2253.02 | 2131.84 | 2120.40 | **5.08** |
| | ***fha*** | A1552VC_A02897 | VCA0112 | 1457.04 | 1409.29 | 1268.37 | 1378.23 | 6962.65 | 8423.01 | 5706.55 | 7030.74 | **5.10** |
| | ***vasD*** | A1552VC_A02898 | VCA0113 | 907.57 | 828.14 | 795.26 | 843.66 | 4013.28 | 5250.54 | 3741.62 | 4335.15 | **5.14** |
| | ***vasE*** | A1552VC_A02899 | VCA0114 | 1539.05 | 1467.41 | 1511.68 | 1506.04 | 7688.85 | 8453.72 | 6328.76 | 7490.45 | **4.97** |
| | ***vasF*** | A1552VC_A02900 | VCA0115 | 936.28 | 740.97 | 835.81 | 837.69 | 4349.21 | 4681.00 | 3674.86 | 4235.02 | **5.06** |
| | ***clpV*** | A1552VC_A02901 | VCA0116 | 2714.52 | 2538.57 | 2747.00 | 2666.70 | 12342.80 | 14252.39 | 11757.13 | 12784.11 | **4.79** |
| | ***vasH*** | A1552VC_A02902 | VCA0117 | 1849.32 | 1569.11 | 1761.74 | 1726.72 | 7548.96 | 8376.48 | 7187.44 | 7704.29 | **4.46** |
| | ***vasI*** | A1552VC_A02903 | VCA0118 | 262.43 | 240.38 | 274.85 | 259.22 | 997.97 | 1115.81 | 974.58 | 1029.46 | **3.97** |
| | ***vasJ*** | A1552VC_A02904 | VCA0119 | 1003.25 | 983.99 | 1169.99 | 1052.41 | 3910.81 | 5173.30 | 4540.02 | 4541.38 | **4.32** |
| | ***vasK*** | A1552VC_A02905 | VCA0120 | 5357.96 | 5123.37 | 5154.57 | 5211.97 | 18884.87 | 26704.97 | 21837.71 | 22475.85 | **4.31** |
| | ***vasL*** | A1552VC_A02906 | VCA0121 | 2165.05 | 1330.04 | 1313.42 | 1602.84 | 6298.82 | 6759.07 | 5504.40 | 6187.43 | **3.86** |
| | ***vasM*** | not annotated | VCA0122 | - | - | - | - | - | - | - | - | **-** |
| | ***vgrG3*** | A1552VC_A02907 | VCA0123 | 4507.79 | 3633.51 | 3470.17 | 3870.49 | 11641.54 | 17214.54 | 12325.56 | 13727.21 | **3.55** |
| | ***tsiV3*** | A1552VC_A02908 | VCA0124 | 1973.70 | 1069.85 | 1073.87 | 1372.47 | 3723.69 | 3987.69 | 3313.21 | 3674.86 | **2.68** |
| Auxiliary cluster 3 | ***paar2*** | A1552VC_A03052 | VCA0284 | 974.55 | 458.32 | 584.24 | 672.37 | 1471.12 | 1713.26 | 1913.93 | 1699.44 | **2.53** |
| | ***tseH*** | A1552VC_A03053 | VCA0285 | 626.01 | 315.67 | 376.23 | 439.30 | 1343.70 | 2119.01 | 2337.70 | 1933.47 | **4.40** |
| | ***tsiH*** | A1552VC_A03054 | VCA0286 | 411.41 | 249.63 | 319.16 | 326.73 | 766.30 | 1007.86 | 1226.81 | 1000.32 | **3.06** |

* according to Matthey, Dresbes Dörr *et al.*, 2018 [20] - genome accession numbers CP028894 and CP028895

[#] according to Heidelberg *et al.*, 2000 [30] - genome accession numbers NC_002505 and NC_002506

**Table S2.** RNA-seq expression data for the T6SS gene cluster in WT and SNP45-converted strain ATCC25872.

| cluster names / comments | gene names | ID (locus tag) in strain ATCC27872[*] | Newly assigned locus tag in strain ATCC27872[$] | Homologous locus tag in strain N16961[#] | ATCC25872 (Exp1) | ATCC25872 (Exp2) | ATCC25872 (Exp1) | AVG - ATCC25872 | ATCC25872-SNP45 converted (Exp1) | ATCC25872-SNP45 converted (Exp2) | ATCC25872-SNP45 converted (Exp3) | AVG - ATCC25872-SNP45 converted | Fold induction [ATCC25872-SNP45 converted / ATCC25872] |
|---|---|---|---|---|---|---|---|---|---|---|---|---|---|
| major T6SS regulators | ***tfoX*** | AT72VC_01697 | GTH07_08410 | VC1153 | 5746.36 | 4043.40 | 5322.23 | 5037.33 | 4000.35 | 3001.05 | 4385.87 | 3795.76 | **0.75** |
| | ***qstR*** | AT72VC_02419 | GTH07_11940 | VC0396 | 165.22 | 245.69 | 257.30 | 222.74 | 256.06 | 216.54 | 243.34 | 238.65 | **1.07** |
| | ***tfoY*** | AT72VC_01140 | GTH07_05665 | VC1722 | 5202.51 | 5065.44 | 5242.38 | 5170.11 | 6040.69 | 5384.32 | 4998.28 | 5474.43 | **1.06** |
| Auxiliary cluster 1 | ***hcp1*** | AT72VC_01454 | GTH07_07215 | VC1415 | 19554.14 | 19652.54 | 18391.42 | 19199.37 | 684.38 | 509.52 | 569.34 | 587.74 | **0.03** |
| | ***vgrG1*** | AT72VC_01453 | GTH07_07210 | VC1416 | 5045.15 | 5650.89 | 5372.93 | 5356.33 | 486.51 | 571.30 | 484.34 | 514.05 | **0.10** |
| | ***tap1*** | AT72VC_01452 | GTH07_07205 | VC1417 | 3486.37 | 3724.64 | 3039.46 | 3416.82 | 423.66 | 503.78 | 385.38 | 437.61 | **0.13** |
| | ***tseL*** | AT72VC_01451 | GTH07_07200 | VC1418 | 11535.01 | 10718.16 | 8489.71 | 10247.63 | 1822.68 | 1887.12 | 1532.20 | 1747.33 | **0.17** |
| | ***tsiV1*** | AT72VC_01450 | GTH07_07195 | VC1419 | 2237.37 | 2221.27 | 1737.74 | 2065.46 | 434.14 | 364.94 | 303.88 | 367.65 | **0.18** |
| Auxiliary cluster 2 | ***hcp2*** | AT72VC_A00658 | GTH07_13450 | VCA0017 | 34594.21 | 31399.13 | 28601.12 | 31531.49 | 1070.80 | 894.84 | 908.14 | 957.93 | **0.03** |
| | ***vgrG2*** | AT72VC_A00659 | GTH07_13455 | VCA0018 | 1691.55 | 1789.25 | 2041.94 | 1840.92 | 197.86 | 164.32 | 140.88 | 167.69 | **0.09** |
| | ***vasW*** | AT72VC_A00660 | GTH07_13460 | VCA0019 | 2132.14 | 2152.76 | 1951.95 | 2078.95 | 268.86 | 205.08 | 174.64 | 216.20 | **0.10** |
| | ***vasX*** | AT72VC_A00661 | GTH07_13465 | VCA0020 | 13234.43 | 12175.86 | 9992.97 | 11801.09 | 1497.95 | 1267.42 | 1102.58 | 1289.32 | **0.11** |
| | ***tsiV2*** | AT72VC_A00662 | GTH07_13470 | VCA0021 | 3450.96 | 2861.52 | 2849.34 | 3053.94 | 600.58 | 518.43 | 468.04 | 529.02 | **0.17** |
| Large/major cluster | ***paar1*** | AT72VC_A00737 | GTH07_13830 | VCA0105 | 384.53 | 358.95 | 385.32 | 376.27 | 290.98 | 307.62 | 292.24 | 296.94 | **0.79** |
| | ***VCA0106*** | AT72VC_A00738 | GTH07_13835 (diff. start codon) | VCA0106 | 706.12 | 782.74 | 846.69 | 778.52 | 484.19 | 574.48 | 548.38 | 535.68 | **0.69** |
| | ***vipA*** | AT72VC_A00739 | GTH07_13840 | VCA0107 | 1544.03 | 3053.33 | 2929.19 | 2508.85 | 442.29 | 544.55 | 525.09 | 503.97 | **0.20** |
| | ***vipB*** | AT72VC_A00740 | GTH07_13845 | VCA0108 | 8279.75 | 12722.05 | 10783.89 | 10595.23 | 1395.53 | 2029.79 | 1857.04 | 1760.78 | **0.17** |
| | ***hsiF*** | AT72VC_A00741 | GTH07_13850 | VCA0109 | 2259.01 | 2625.88 | 2116.72 | 2333.87 | 246.75 | 345.20 | 331.82 | 307.92 | **0.13** |
| | ***vasA*** | AT72VC_A00742 | GTH07_13855 | VCA0110 | 2802.86 | 3578.50 | 2946.94 | 3109.43 | 366.63 | 570.66 | 497.15 | 478.15 | **0.15** |
| | ***vasB*** | AT72VC_A00743 | GTH07_13860 | VCA0111 | 733.66 | 652.13 | 803.59 | 729.80 | 97.77 | 135.66 | 140.88 | 124.77 | **0.17** |
| | ***fha*** | AT72VC_A00744 | GTH07_13865 | VCA0112 | 2701.57 | 2296.16 | 1802.38 | 2266.70 | 342.19 | 445.83 | 376.06 | 388.03 | **0.17** |
| | ***vasD*** | AT72VC_A00745 | GTH07_13870 | VCA0113 | 1634.51 | 1662.30 | 1131.88 | 1476.23 | 251.40 | 259.22 | 222.38 | 244.33 | **0.17** |
| | ***vasE*** | AT72VC_A00746 | GTH07_13875 | VCA0114 | 2605.19 | 2778.41 | 2040.67 | 2474.76 | 425.99 | 527.35 | 437.77 | 463.70 | **0.19** |
| | ***vasF*** | AT72VC_A00747 | GTH07_13880 | VCA0115 | 1576.49 | 1380.07 | 1097.65 | 1351.40 | 235.11 | 280.23 | 223.54 | 246.30 | **0.18** |
| | ***clpV*** | AT72VC_A00748 | GTH07_13885 | VCA0116 | 4491.46 | 4472.67 | 3551.53 | 4171.89 | 890.39 | 1061.07 | 867.39 | 939.62 | **0.23** |
| | ***vasH*** | AT72VC_A00749 | GTH07_13890 | VCA0117 | 3205.10 | 3151.97 | 2592.04 | 2983.03 | 665.76 | 737.53 | 649.67 | 684.32 | **0.23** |
| | ***vasI*** | AT72VC_A00750 | GTH07_13895 | VCA0118 | 432.72 | 396.39 | 394.19 | 407.77 | 83.80 | 120.37 | 121.09 | 108.42 | **0.27** |
| | ***vasJ*** | AT72VC_A00751 | GTH07_13900 | VCA0119 | 1448.64 | 1455.88 | 1339.75 | 1414.75 | 301.45 | 376.41 | 341.14 | 339.66 | **0.24** |
| | ***vasK*** | AT72VC_A00752 | GTH07_13905 | VCA0120 | 10261.43 | 10150.05 | 8146.22 | 9519.23 | 1559.64 | 2015.78 | 1793.00 | 1789.47 | **0.19** |
| | ***vasL*** | AT72VC_A00753 | GTH07_13910 | VCA0121 | 3757.80 | 2878.88 | 2001.38 | 2879.35 | 523.76 | 552.83 | 472.70 | 516.43 | **0.18** |
| | ***vasM*** | not annotated | not annotated | VCA0122 | - | - | - | - | - | - | - | - | **-** |
| | ***vgrG3*** | AT72VC_A00754 | GTH07_13915 | VCA0123 | 7884.40 | 6577.94 | 5367.86 | 6610.07 | 1783.11 | 1917.06 | 1458.85 | 1719.67 | **0.26** |
| | ***tsiV3*** | AT72VC_A00755 | GTH07_13920 | VCA0124 | 2256.06 | 1586.49 | 1221.87 | 1688.14 | 669.25 | 664.28 | 561.19 | 631.57 | **0.37** |
| Auxiliary cluster 3 | ***paar2*** | AT72VC_A00904 | GTH07_14650 | VCA0284 | 973.63 | 926.14 | 866.97 | 922.24 | 684.38 | 644.54 | 470.37 | 599.76 | **0.65** |
| | ***tseH*** | AT72VC_A00905 | GTH07_14655 | VCA0285 | 765.13 | 906.96 | 788.38 | 820.16 | 543.55 | 445.83 | 284.09 | 424.49 | **0.52** |
| | ***tsiH*** | AT72VC_A00906 | GTH07_14660 (diff. start codon) | VCA0286 | 472.06 | 569.02 | 487.99 | 509.69 | 385.25 | 277.05 | 224.71 | 295.67 | **0.58** |

* according to Prokka annotation used for RNA-seq read mapping in this work

[$] according to NCBI's PGAP annotation - genome accession numbers CP047305 and CP047306.

[#] according to Heidelberg *et al.*, 2000 [30] - genome accession numbers NC_002505 and NC_002506

**Table S3.** *Vibrio cholerae* and *Escherichia coli* strains and plasmids used in this study.

| Strain names | Genotype / description* | Internal strain number | Reference |
|---|---|---|---|
| ***V. cholerae*** | | | |
| A1552 | Wild-type, O1 El Tor Inaba, isolated in 1991 in Peru; $Rif^R$. | MB_1 | [19]; [20] |
| ATCC25872 | Wild-type, non-O1 strain (O37); isolated in 1965 in Czechoslovakia; intermediate $Strep^R$ | MB_276 | [21] |
| A1552-*aph*#1 | A1552 with *aph* cassette in position #1 on chr 1; $Rif^R$, $Kan^R$ | MB_6911 | This study |
| A1552-*aph*#2 | A1552 with *aph* cassette in position #2 on chr 1; $Rif^R$, $Kan^R$ | MB_6912 | This study |
| A1552-*aph*#3 | A1552 with *aph* cassette in position #3 on chr 1; $Rif^R$, $Kan^R$ | MB_6913 | This study |
| A1552-*aph*#4 | A1552 with *aph* cassette in position #4 on chr 1; $Rif^R$, $Kan^R$ | MB_6914 | This study |
| A1552-*aph*#5 | A1552 with *aph* cassette in position #5 on chr 1; $Rif^R$, $Kan^R$ | MB_6915 | This study |
| A1552-*aph*#6 | A1552 with *aph* cassette in position #6 on chr 1; $Rif^R$, $Kan^R$ | MB_6916 | This study |
| A1552-*aph*#7 | A1552 with *aph* cassette in position #7 on chr 1; $Rif^R$, $Kan^R$ | MB_6917 | This study |
| A1552-*aph*#8 | A1552 with *aph* cassette in position #8 on chr 1; $Rif^R$, $Kan^R$ | MB_6918 | This study |
| A1552-*aph*#9 | A1552 with *aph* cassette in position #9 on chr 1; $Rif^R$, $Kan^R$ | MB_6919 | This study |
| A1552-*aph*#10 | A1552 with *aph* cassette in position #10 on chr 1; $Rif^R$, $Kan^R$ | MB_6920 | This study |
| A1552-*aph*#11 | A1552 with *aph* cassette in position #11 on chr 1; $Rif^R$, $Kan^R$ | MB_6921 | This study |
| A1552-*aph*#12 | A1552 with *aph* cassette in position #12 on chr 1; $Rif^R$, $Kan^R$ | MB_6922 | This study |
| A1552-*aph*#13 | A1552 with *aph* cassette in position #13 on chr 1; $Rif^R$, $Kan^R$ | MB_6923 | This study |
| A1552-*aph*#14 | A1552 with *aph* cassette in position #14 on chr 1; $Rif^R$, $Kan^R$ | MB_6924 | This study |
| A1552-*aph*#15 | A1552 with *aph* cassette in position #15 on chr 1; $Rif^R$, $Kan^R$ | MB_6925 | This study |
| A1552-*aph*#16 | A1552 with *aph* cassette in position #16 on chr 1; $Rif^R$, $Kan^R$ | MB_6926 | This study |
| A1552-*aph*#17 | A1552 with *aph* cassette in position #17 on chr 1; $Rif^R$, $Kan^R$ | MB_6927 | This study |
| A1552-*aph*#18 | A1552 with *aph* cassette in position #18 on chr 1; $Rif^R$, $Kan^R$ | MB_6928 | This study |
| A1552-*aph*#19 | A1552 with *aph* cassette in position #19 on chr 1; $Rif^R$, $Kan^R$ | MB_6929 | This study |
| A1552-*aph*#20 | A1552 with *aph* cassette in position #20 on chr 1; $Rif^R$, $Kan^R$ | MB_6930 | This study |
| A1552-*aph*#21 | A1552 with *aph* cassette in position #21 on chr 1; $Rif^R$, $Kan^R$ | MB_6931 | This study |
| A1552-*aph*#22 | A1552 with *aph* cassette in position #22 on chr 1; $Rif^R$, $Kan^R$ | MB_6932 | This study |
| A1552-*aph*#23 | A1552 with *aph* cassette in position #23 on chr 1; $Rif^R$, $Kan^R$ | MB_6933 | This study |
| A1552-*aph*#24 | A1552 with *aph* cassette in position #24 on chr 1; $Rif^R$, $Kan^R$ | MB_6934 | This study |
| A1552-*aph*#25 | A1552 with *aph* cassette in position #25 on chr 1; $Rif^R$, $Kan^R$ | MB_6935 | This study |
| A1552-*aph*#26 | A1552 with *aph* cassette in position #26 on chr 1; $Rif^R$, $Kan^R$ | MB_6936 | This study |
| A1552-*aph*#27 | A1552 with *aph* cassette in position #27 on chr 1; $Rif^R$, $Kan^R$ | MB_6937 | This study |
| A1552-*aph*#28 | A1552 with *aph* cassette in position #28 on chr 1; $Rif^R$, $Kan^R$ | MB_6938 | This study |
| A1552-*aph*#29 | A1552 with *aph* cassette in position #29 on chr 1; $Rif^R$, $Kan^R$ | MB_6939 | This study |
| A1552-*aph*#30 | A1552 with *aph* cassette in position #30 on chr 1; $Rif^R$, $Kan^R$ | MB_6940 | This study |
| A1552-*aph*#31 | A1552 with *aph* cassette in position #31 on chr 2; $Rif^R$, $Kan^R$ | MB_6941 | This study |
| A1552-*aph*#32 | A1552 with *aph* cassette in position #32 (15kb upstream of T6SS large cluster) on chr 2; $Rif^R$, $Kan^R$ | MB_6942 | This study |
| A1552-*aph*#33 | A1552 with *aph* cassette in position #33 on chr 2; $Rif^R$, $Kan^R$ | MB_6943 | This study |
| A1552-*aph*#34 | A1552 with *aph* cassette in position #34 on chr 2; $Rif^R$, $Kan^R$ | MB_6944 | This study |
| A1552-*aph*#35 | A1552 with *aph* cassette in position #35 on chr 2; $Rif^R$, $Kan^R$ | MB_6945 | This study |
| A1552-*aph*#36 | A1552 with *aph* cassette in position #36 on chr 2; $Rif^R$, $Kan^R$ | MB_6946 | This study |
| A1552-*aph*#37 | A1552 with *aph* cassette in position #37 on chr 2; $Rif^R$, $Kan^R$ | MB_6947 | This study |
| A1552-*aph*#38 | A1552 with *aph* cassette in position #38 on chr 2; $Rif^R$, $Kan^R$ | MB_6948 | This study |
| A1552-*aph*#39 | A1552 with *aph* cassette in position #39 on chr 2; $Rif^R$, $Kan^R$ | MB_6949 | This study |

| | | | |
|---|---|---|---|
| A1552-*aph*#40 | A1552 with *aph* cassette in position #40 on chr 2; $Rif^R$, $Kan^R$ | MB_6950 | This study |
| A1552-*aph*#42 | A1552 with *aph* cassette in position #42 on chr 2 (upstream of *paar1/VCA0105*); $Rif^R$, $Kan^R$ | MB_7922 | This study |
| ATCC25872-*aph*#1$^{A1552}$ hybrid clones 1-20 | ATCC25872 transformed with gDNA of strain A1552-*aph*#1 (#6911). 20 hybrid transformants selected; intermediate $Strep^R$, $Kan^R$ | MB_7001 to 7020 | This study |
| ATCC25872-*aph*#2$^{A1552}$ hybrid clones 1-20 | ATCC25872 transformed with gDNA of strain A1552-*aph*#2 (#6912). 20 hybrid clones selected; intermediate $Strep^R$, $Kan^R$ | MB_7021 to 7040 | This study |
| ATCC25872-*aph*#3$^{A1552}$ hybrid clones 1-20 | ATCC25872 transformed with gDNA of strain A1552-*aph*#3 (#6913). 20 hybrid clones selected; intermediate $Strep^R$, $Kan^R$ | MB_7041 to 7060 | This study |
| ATCC25872-*aph*#4$^{A1552}$ hybrid clones 1-20 | ATCC25872 transformed with gDNA of strain A1552-*aph*#4 (#6914). 20 hybrid clones selected; intermediate $Strep^R$, $Kan^R$ | MB_7061 to 7080 | This study |
| ATCC25872-*aph*#5$^{A1552}$ hybrid clones 1-20 | ATCC25872 transformed with gDNA of strain A1552-*aph*#5 (#6915). 20 hybrid clones selected; intermediate $Strep^R$, $Kan^R$ | MB_7081 to 7100 | This study |
| ATCC25872-*aph*#6$^{A1552}$ hybrid clones 1-20 | ATCC25872 transformed with gDNA of strain A1552-*aph*#6 (#6916). 20 hybrid clones selected; intermediate $Strep^R$, $Kan^R$ | MB_7101 to 7120 | This study |
| ATCC25872-*aph*#7$^{A1552}$ hybrid clones 1-20 | ATCC25872 transformed with gDNA of strain A1552-*aph*#7 (#6917). 20 hybrid clones selected; intermediate $Strep^R$, $Kan^R$ | MB_7121 to 7140 | This study |
| ATCC25872-*aph*#8$^{A1552}$ hybrid clones 1-20 | ATCC25872 transformed with gDNA of strain A1552-*aph*#8 (#6918). 20 hybrid clones selected intermediate $Strep^R$, $Kan^R$ | MB_7141 to 7160 | This study |
| ATCC25872-*aph*#9$^{A1552}$ hybrid clones 1-20 | ATCC25872 transformed with gDNA of strain A1552-*aph*#9 (#6919). 20 hybrid clones selected; intermediate $Strep^R$, $Kan^R$ | MB_7161 to 7180 | This study |
| ATCC25872-*aph*#10$^{A1552}$ hybrid clones 1-20 | ATCC25872 transformed with gDNA of strain A1552-*aph*#10 (#6920). 20 hybrid clones selected; intermediate $Strep^R$, $Kan^R$ | MB_7181 to 7200 | This study |
| ATCC25872-*aph*#11$^{A1552}$ hybrid clones 1-20 | ATCC25872 transformed with gDNA of strain A1552-*aph*#11 (#6921). 20 hybrid clones selected; intermediate $Strep^R$, $Kan^R$ | MB_7201 to 7220 | This study |
| ATCC25872-*aph*#12$^{A1552}$ hybrid clones 1-20 | ATCC25872 transformed with gDNA of strain A1552-*aph*#12 (#6922). 20 hybrid clones selected; intermediate $Strep^R$, $Kan^R$ | MB_7221 to 7240 | This study |
| ATCC25872-*aph*#13$^{A1552}$ hybrid clones 1-20 | ATCC25872 transformed with gDNA of strain A1552-*aph*#13 (#6923). 20 hybrid clones selected; intermediate $Strep^R$, $Kan^R$ | MB_7241 to 7260 | This study |
| ATCC25872-*aph*#14$^{A1552}$ hybrid clones 1-20 | ATCC25872 transformed with gDNA of strain A1552-*aph*#14 (#6924). 20 hybrid clones selected; intermediate $Strep^R$, $Kan^R$ | MB_7261 to 7280 | This study |

| | | | |
|---|---|---|---|
| ATCC25872-*aph*#15$^{A1552}$ hybrid clones 1-20 | ATCC25872 transformed with gDNA of strain A1552-*aph*#15 (#6925). 20 hybrid clones selected; intermediate $Strep^R$, $Kan^R$ | MB_7281 to 7300 | This study |
| ATCC25872-*aph*#16$^{A1552}$ hybrid clones 1-20 | ATCC25872 transformed with gDNA of strain A1552-*aph*#16 (#6926). 20 hybrid clones selected; intermediate $Strep^R$, $Kan^R$ | MB_7301 to 7320 | This study |
| ATCC25872-*aph*#17$^{A1552}$ hybrid clones 1-20 | ATCC25872 transformed with gDNA of strain A1552-*aph*#17 (#6927). 20 hybrid clones selected; intermediate $Strep^R$, $Kan^R$ | MB_7321 to 7340 | This study |
| ATCC25872-*aph*#18$^{A1552}$ hybrid clones 1-20 | ATCC25872 transformed with gDNA of strain A1552-*aph*#18 (#6928). 20 hybrid clones selected; intermediate $Strep^R$, $Kan^R$ | MB_7341 to 7360 | This study |
| ATCC25872-*aph*#19$^{A1552}$ hybrid clones 1-20 | ATCC25872 transformed with gDNA of strain A1552-*aph*#19 (#6929). 20 hybrid clones selected; intermediate $Strep^R$, $Kan^R$ | MB_7361 to 7380 | This study |
| ATCC25872-*aph*#20$^{A1552}$ hybrid clones 1-20 | ATCC25872 transformed with gDNA of strain A1552-*aph*#20 (#6930). 20 hybrid clones selected; intermediate $Strep^R$, $Kan^R$ | MB_7381 to 7400 | This study |
| ATCC25872-*aph*#21$^{A1552}$ hybrid clones 1-20 | ATCC25872 transformed with gDNA of strain A1552-*aph*#21 (#6931). 20 hybrid clones selected; intermediate $Strep^R$, $Kan^R$ | MB_7401 to 7420 | This study |
| ATCC25872-*aph*#22$^{A1552}$ hybrid clones 1-20 | ATCC25872 transformed with gDNA of strain A1552-*aph*#22 (#6932). 20 hybrid clones selected; intermediate $Strep^R$, $Kan^R$ | MB_7421 to 7440 | This study |
| ATCC25872-*aph*#23$^{A1552}$ hybrid clones 1-20 | ATCC25872 transformed with gDNA of strain A1552-*aph*#23 (#6933). 20 hybrid clones selected; intermediate $Strep^R$, $Kan^R$ | MB_7441 to 7460 | This study |
| ATCC25872-*aph*#24$^{A1552}$ hybrid clones 1-20 | ATCC25872 transformed with gDNA of strain A1552-*aph*#24 (#6934). 20 hybrid clones selected; intermediate $Strep^R$, $Kan^R$ | MB_7461 to 7480 | This study |
| ATCC25872-*aph*#25$^{A1552}$ hybrid clones 1-20 | ATCC25872 transformed with gDNA of strain A1552-*aph*#25 (#6935). 20 hybrid clones selected; intermediate $Strep^R$, $Kan^R$ | MB_7481 to 7500 | This study |
| ATCC25872-*aph*#26$^{A1552}$ hybrid clones 1-20 | ATCC25872 transformed with gDNA of strain A1552-*aph*#26 (#6936). 20 hybrid clones selected; intermediate $Strep^R$, $Kan^R$ | MB_7501 to 7520 | This study |
| ATCC25872-*aph*#27(A1552) hybrid clones 1-20 | ATCC25872 transformed with gDNA of strain A1552-*aph*#27 (#6937). 20 hybrid clones selected; intermediate $Strep^R$, $Kan^R$ | MB_7521 to 7540 | This study |
| ATCC25872-*aph*#28$^{A1552}$ hybrid clones 1-20 | ATCC25872 transformed with gDNA of strain A1552-*aph*#28 (#6938). 20 hybrid clones selected; intermediate $Strep^R$, $Kan^R$ | MB_7541 to 7560 | This study |
| ATCC25872-*aph*#29$^{A1552}$ hybrid clones 1-20 | ATCC25872 transformed with gDNA of strain A1552-*aph*#29 (#6939). 20 hybrid clones selected; intermediate $Strep^R$, $Kan^R$ | MB_7561 to 7580 | This study |

| | | | |
|---|---|---|---|
| ATCC25872-*aph*#30$^{A1552}$ hybrid clones 1-20 | ATCC25872 transformed with gDNA of strain A1552-*aph*#30 (#6940). 20 hybrid clones selected; intermediate Strep$^{R}$, Kan$^{R}$ | MB_7581 to 7600 | This study |
| ATCC25872-*aph*#31$^{A1552}$ hybrid clones 1-20 | ATCC25872 transformed with gDNA of strain A1552-*aph*#31 (#6941). 20 hybrid clones selected; intermediate Strep$^{R}$, Kan$^{R}$ | MB_7601 to 7620 | This study |
| ATCC25872-*aph*#32$^{A1552}$ hybrid clones 1-20 | ATCC25872 transformed with gDNA of strain A1552-*aph*#32 (#6942). 20 hybrid clones selected; intermediate Strep$^{R}$, Kan$^{R}$ | MB_7621 to 7640 | This study |
| ATCC25872-*aph*#33$^{A1552}$ hybrid clones 1-20 | ATCC25872 transformed with gDNA of strain A1552-*aph*#33 (#6943). 20 hybrid clones selected; intermediate Strep$^{R}$, Kan$^{R}$ | MB_7641 to 7660 | This study |
| ATCC25872-*aph*#34$^{A1552}$ hybrid clones 1-20 | ATCC25872 transformed with gDNA of strain A1552-*aph*#34 (#6944). 20 hybrid clones selected; intermediate Strep$^{R}$, Kan$^{R}$ | MB_7661 to 7680 | This study |
| ATCC25872-*aph*#35$^{A1552}$ hybrid clones 1-20 | ATCC25872 transformed with gDNA of strain A1552-*aph*#35 (#6945). 20 hybrid clones selected; intermediate Strep$^{R}$, Kan$^{R}$ | MB_7681 to 7700 | This study |
| ATCC25872-*aph*#36$^{A1552}$ hybrid clones 1-20 | ATCC25872 transformed with gDNA of strain A1552-*aph*#36 (#6946). 20 hybrid clones selected; intermediate Strep$^{R}$, Kan$^{R}$ | MB_7701 to 7720 | This study |
| ATCC25872-*aph*#37$^{A1552}$ hybrid clones 1-20 | ATCC25872 transformed with gDNA of strain A1552-*aph*#37 (#6947). 20 hybrid clones selected; intermediate Strep$^{R}$, Kan$^{R}$ | MB_7721 to 7740 | This study |
| ATCC25872-*aph*#38$^{A1552}$ hybrid clones 1-20 | ATCC25872 transformed with gDNA of strain A1552-*aph*#38 (#6948). 20 hybrid clones selected; intermediate Strep$^{R}$, Kan$^{R}$ | MB_7741 to 7760 | This study |
| ATCC25872-*aph*#39$^{A1552}$ hybrid clones 1-20 | ATCC25872 transformed with gDNA of strain A1552-*aph*#39 (#6949). 20 hybrid clones selected; intermediate Strep$^{R}$, Kan$^{R}$ | MB_7761 to 7780 | This study |
| ATCC25872-*aph*#40$^{A1552}$ hybrid clones 1-20 | ATCC25872 transformed with gDNA of strain A1552-*aph*#40 (#6950). 20 hybrid clones selected; intermediate Strep$^{R}$, Kan$^{R}$ | MB_7781 to 7800 | This study |
| ATCC25872-*aph*#42$^{A1552}$ hybrid clones 1-20 | ATCC25872 transformed with gDNA of strain A1552-*aph*#42 (#7922). 20 hybrid clones selected; intermediate Strep$^{R}$, Kan$^{R}$ | MB_7951 to 7970 | This study |
| ATCC25872-*aph*#32 | ATCC25872 with *aph* cassette in position #32 on chr 2 (15kb upstream of T6SS large cluster); intermediate Strep$^{R}$, Kan$^{R}$ | MB_6982 | This study |
| A1552-*aph*#32$^{ATCC25872}$ hybrid clones 1-20 | A1552 transformed with gDNA of strain ATCC25872-*aph*#32 (#6982). 20 hybrid clones selected; Rif$^{R}$, Kan$^{R}$ | ND_178 to 197 | This study |
| ATCC25872-*aph*#42 | ATCC25872 with *aph* cassette in position #42 on chr 2 (upstream of *paar1/VCA0105*); intermediate Strep$^{R}$, Kan$^{R}$ | MB_7995 | This study |
| A1552-*aph*#42$^{ATCC25872}$ hybrid clones | A1552 transformed with gDNA of strain ATCC25872-*aph*#42 (#7995). 20 hybrid clones selected; Rif$^{R}$, Kan$^{R}$ | ND_228 to 247 | This study |

| | | | |
|---|---|---|---|
| 1-20 | | | |
| A1552-T6SS [SNP45-T] | A1552 with SNP45 in intergenic region between *VCA0106* and *VCA0107* (*vipA*) converted from G to T by site-directed mutation; $Rif^R$ | MB_9063 | This study |
| A1552-T6SS [SNP45-C] | A1552 with SNP45 in intergenic region between *VCA0106* and *VCA0107* (*vipA*) converted from G to C by site-directed mutation; $Rif^R$ | ND_333 | This study |
| A1552-T6SS [SNP45-A] | A1552 with SNP45 in intergenic region between *VCA0106* and *VCA0107* (*vipA*) converted from G to A by site-directed mutation; $Rif^R$ | ND_334 | This study |
| ATCC25872-T6SS [SNP45-G] | ATCC25872 with SNP45 in intergenic region between *VCA0106* and *VCA0107* (*vipA*) converted from T to G by site-directed mutation; intermediate $Strep^R$ | MB_9168 | This study |
| A1552-*vipA-sfGFP*v2 | A1552 carrying *vipA-sfGFP* translational fusion (version 2; without ATG at start of *sfGFP*); TransFLP method; $Rif^R$ | MB_3909 | [18] |
| A1552-T6SS [SNP45-T]-*vipA-sfGFPv2* | A1552-T6SS[SNP45-T] carrying *vipA-sfGFP* translational fusion (v2: without ATG at start of *sfGFP*), TransFLP method; $Rif^R$ | ND_273 | This study |
| ATCC25872-*vipA-sfGFP*v2 | ATCC25872 carrying *vipA-sfGFP* translational fusion (version 2; without ATG at start of *sfGFP*); TransFLP method; intermediate $Strep^R$ | ND_302 | This study |
| ATCC25872- *vipA-sfGFP*v2 -T6SS [SNP45-G] | ATCC25872-*vipA-sfGFP*v2 with SNP45 in intergenic region between *VCA0106* and *VCA0107* (*vipA*) converted from T to G by site-directed mutation; intermediate $Strep^R$ | ND_643 | This study |
| A1552Δ*tfoX* | A1552 with *tfoX* (*VC1153*) deleted; $Rif^R$ | MB_45 | [1] |
| A1552Δ*qstR* | A1552 with *qstR* (*VC0396*) deleted; $Rif^R$. | MB_600 | [22] |
| A1552Δ*tfoY* | A1552 with *tfoY* (*VC1722*) deleted; $Rif^R$ | MB_828 | [18] |
| A1552Δ*vipB* | A1552 with *vipB* (*VCA0108*) deleted; $Rif^R$ | MB_598 | [4] |
| A1552Δ*vasK* | A1552 with *vasK* (*VCA0120*) deleted; $Rif^R$ | MB_585 | [4] |
| A1552Δ*hapR* | A1552 with *hapR* (*VC0583*) deleted; $Rif^R$ | MB_3 | [1] |
| A1552-T6SS [SNP45-T] Δ*tfoX* | A1552-T6SS[SNP45-T] with *tfoX* (*VC1153*) deleted; $Rif^R$ | ND_320 | This study |
| A1552-T6SS [SNP45-T] Δ*qstR* | A1552-T6SS[SNP45-T] with *qstR* (*VC0396*) deleted; $Rif^R$ | ND_322 | This study |
| A1552-T6SS [SNP45-T] Δ*tfoY* | A1552-T6SS[SNP45-T] with *tfoY* (*VC1722*) deleted; $Rif^R$ | ND_321 | This study |
| A1552-T6SS [SNP45-T] Δ*vipB* | A1552-T6SS[SNP45-T] with *vipB* (*VCA0108*) deleted; $Rif^R$ | ND_339 | This study |
| A1552-T6SS [SNP45-T] Δ*vasK* | A1552-T6SS[SNP45-T] with *vasK* (*VCA0120*) deleted; $Rif^R$ | ND_323 | This study |
| A1552-T6SS [SNP45-T] Δ*hapR* | A1552-T6SS[SNP45-T] with *hapR* (*VC0583*) deleted; $Rif^R$ | ND_324 | This study |
| A1552ΔIGR-ALL | A1552 with deletion of the intergenic region (396 bp) between *VCA0106* and *VCA0107* (*vipA*); 10 bp downstream of *VCA0106* and 25 bp upstream of *vipA* were kept; $Rif^R$ | ND_370 | This study |
| A1552ΔIGR276 | A1552 with deletion of 276 bp in intergenic region between *VCA0106* and *VCA0107* (*vipA*); $Rif^R$ | ND_567 | This study |
| A1552ΔIGR336 | A1552 with deletion of 336 bp in intergenic region between *VCA0106* and *VCA0107* (*vipA*); $Rif^R$ | ND_569 | This study |
| A1552-T6SS [SNP45-T] ΔIGR276 | A1552-T6SS[SNP45-T] with deletion of 276 bp in intergenic region between *VCA0106* and *VCA0107* (*vipA*); $Rif^R$ | ND_568 | This study |
| A1552-T6SS [SNP45-T] ΔIGR336 | A1552-T6SS[SNP45-T] with deletion of 336 bp in intergenic region between *VCA0106* and *VCA0107* (*vipA*); $Rif^R$ | ND_570 | This study |
| A1552-T6SS [SNP45P-T] ΔIGR60 | A1552-T6SS[SNP45-T] with deletion of 60 bp in intergenic region between *VCA0106* and *VCA0107* (*vipA*); $Rif^R$ | ND_755 | This study |

| | | | |
|---|---|---|---|
| W10G | Environmental isolate (non-O1/non-O139) collected in 2004 in Waddell Creek (CA, USA) | MB_5537 | [23] |
| SA3G | Environmental isolate (non-O1/non-O139) collected in 2004 in Old Salinas River (CA, USA) | MB_957 | [23] |
| SA5Y | Environmental isolate (non-O1/non-O139) collected in 2004 in Old Salinas River (CA, USA) | MB_353 | [23] |
| SL4G | Environmental isolate (non-O1/non-O139) collected in 2004 in San Lorenzo River (CA, USA); $Amp^R$ | MB_955 | [23] |
| SL4G-T6SS [SNP45-G] | SL4G with SNP45 in intergenic region between *VCA0106* and *VCA0107* (*vipA*) converted from T to G by site-directed mutation; $Amp^R$ | ND_513 | This study |
| SL5Y | Environmental isolate (non-O1/non-O139) collected in 2004 in San Lorenzo River (CA, USA) | MB_954 | [23] |
| SL5Y-T6SS [SNP45-G] | SL5Y with SNP45 in intergenic region between *VCA0106* and *VCA0107* (*vipA*) converted from T to G by site-directed mutation | ND_512 | This study |
| SO5Y | Environmental isolate (non-O1/non-O139) collected in 2004 in Soquel Creek (CA, USA) | MB_960 | [23] |
| L6G | Wild-type; environmental isolate (non-O1/ non-O139) collected in 2004 in Lagunitas Creek (CA, USA); $Amp^R$. | MB_956 | [23] |
| L6G-T6SS [SNP45-G] | L6G with SNP45 in intergenic region between *VCA0106* and *VCA0107* (*vipA*) converted from T to G by site-directed mutation; $Amp^R$. | ND_509 | This study |
| SL6Y | Wild-type; environmental isolate (non-O1/ non-O139) collected in 2004 in San Lorenzo River (CA, USA) | MB_953 | [23] |
| SL6Y-T6SS [SNP45-G] | SL6Y with SNP45 in intergenic region between *VCA0106* and *VCA0107* (*vipA*) converted from T to G by site-directed mutation | ND_511 | This study |
| SP6G | Wild-type; environmental isolate (non-O1/ non-O139) collected in 2004 in San Pedro Creek (CA, USA). | MB_964 | [23] |
| SP6G-T6SS [SNP45-G] | SP6G with SNP45 in intergenic region between *VCA0106* and *VCA0107* (*vipA*) converted from T to G by site-directed mutation | ND_514 | This study |
| SP7G | Wild-type; environmental isolate (non-O1/ non-O139) collected in 2004 in San Pedro Creek (CA, USA) | MB_952 | [23] |
| W6G | Wild-type; environmental isolate (non-O1/ non-O139) collected in 2004 in Waddell Creek (CA, USA) | MB_354 | [23] |
| W7G | Wild-type; environmental isolate (non-O1/ non-O139) collected in 2004 in Waddell Creek (CA, USA) | MB_962 | [23] |
| E7G | Wild-type; environmental isolate (non-O1/ non-O139) collected in 2004 in Moss Landing Harbor (CA, USA) | MB_963 | [23] |
| SA7G | Wild-type; environmental isolate (non-O1/ non-O139) collected in 2004 in Old Salinas River (CA, USA) | MB_959 | [23] |
| SA10G | Wild-type; environmental isolate (non-O1/ non-O139) collected in 2004 in Old Salinas River (CA, USA) | MB_5539 | [23] |
| C6706 ($Strep^S$) (original) | Wild-type; O1 El Tor Inaba collected in 1991 in Peru. Original isolate before introduction of streptomycin resistance mutation; non-mutated *luxO;* $Strep^S$ | MB_4522 | Gift from J. Mekalanos (Harvard); [24] |
| C6706-T6SS [SNP45-T] | C6706 with SNP45 in intergenic region between *VCA0106* and *VCA0107* (*vipA*) converted from G to T by site-directed mutation; $Strep^S$ | ND_538 | This study |
| E7946 | Wild-type; O1 El Tor Ogawa isolated in 197 in Bahrain. $Strep^R$ | MB_2600 | Lab stock; [25] |
| E7946-T6SS [SNP45-T] | E7946 with SNP45 in intergenic region between *VCA0106* and *VCA0107* (*vipA*) converted from G to T by site-directed mutation; $Strep^R$ | ND_539 | This study |

| | | | |
|---|---|---|---|
| P27459 | Wild-type; O1 El Tor Inaba isolated in 1976 in Bangladesh; $Strep^R$ | MB_1504 | [26] |
| P27459-T6SS [SNP45-T] | P27459 with SNP45 in intergenic region between *VCA0106* and *VCA0107* (*vipA*) converted from G to T by site-directed mutation; $Strep^R$ | ND_540 | This study |
| DRC-193A | Wild-type; O1 isolated in 2011 in the Democratic Republic of Congo; $Strep^R$ | MB_1954 | [4] |
| DRC-193A -T6SS [SNP45-T] | DRC-193A with SNP45 in intergenic region between *VCA0106* and *VCA0107* (*vipA*) converted from G to T by site-directed mutation; $Strep^R$ | ND_541 | This study |
| N16961-*hapR*$^{Rep}$ | Wild-type; O1 El Tor Inaba isolated in 1975 in Bangladesh. *hapR* frameshift mutation repaired; $Strep^R$ | MB_5663 | [24] |
| N16961-*hapR*$^{Rep}$-T6SS [SNP45-T] | N16961-*hapR*$^{Rep}$ with SNP45 in intergenic region between *VCA0106* and *VCA0107* (*vipA*) converted from G to T by site-directed mutation; $Strep^R$ | ND_537 | This study |
| A1552-T6SS [SNP45-T]-chg[-10box] | A1552-T6SS[SNP45-T] with site-directed mutation (AA to GC) in -10 element in intergenic region between *VCA0106* and *VCA0107* (*vipA*); RifR, GentR | ND_757 | This study |
| A1552-Tn*tfoX*-strep (TnTfoX) | A1552 containing mini-Tn7-*araC*-$P_{BAD}$-*tfoX*-strep; RifR, GentR | MB_3420 | [18] |
| A1552-chg[-10box]-Tn*tfoX*-strep (mut-10; TnTfoX) | A1552-Tn*tfoX*-strep with site-directed mutation (AA to GC) in -10 element in intergenic region between *VCA0106* and *VCA0107* (*vipA*); RifR, GentR | ND_768 | This study |
| A1552-Tn*qstR* (TnQstR) | A1552 containing mini-Tn7-*araC*-$P_{BAD}$-*qstR*; RifR, GentR | MB_5501 | [27] |
| A1552-chg[-10box]-Tn*qstR* (mut-10; TnQstR) | A1552-Tn*qstR* with site-directed mutation (AA to GC) in -10 element in intergenic region between *VCA0106* and *VCA0107* (*vipA*); RifR, GentR | ND_770 | This study |
| A1552-Tn*tfoY*-strep (TnTfoY) | A1552 containing mini-Tn7-*araC*-$P_{BAD}$-*tfoY*-strep; RifR, GentR | MB_2978 | [18] |
| A1552-chg[-10box]-Tn*tfoY*-strep (mut-10; TnTfoY) | A1552-Tn*tfoY*-strep with site-directed mutation (AA to GC) in -10 element in intergenic region between *VCA0106* and *VCA0107* (*vipA*); RifR, GentR | ND_769 | This study |
| ***E. coli*** | | | |
| TOP10 | F- mcrA Δ(mrr-hsdRMS-mcrBC) φ80lacZΔM15 ΔlacX74 nupG recA1 araΔ139 Δ(ara-leu)7697 galE15 galK16 rpsL(StrR) endA1λ-. | MB_741 | Invitrogen |
| TOP10-TnKan | TOP10 containing mini-Tn7-*aph* ($Kan^R$); $Strep^R$, $Kan^R$, $Gent^R$. | MB_4119 | [18] |
| TOP10-TnGFP | TOP10 containing mini-Tn7-GFP; $Strep^R$, $Cm^R$, $Gent^R$. | MB_4482 | This study |
| MC4100-TnGFP | MC4100 containing mini-Tn7-GFP; $Strep^R$, $Cm^R$, $Gent^R$. | MB_3930 | This study |
| DH5α | F- endA1 glnV44 thi-1 recA1 relA1 gyrA96 deoR nupG φ80lacZΔM15 Δ(lacZYA-argF) U169 hsdR17 ($rK^-$ $mK^+$) phoA, λ-. | MB_736 | [28] |
| S17-1λpir | Tpr Smr recA thi pro hsdR2M1 RP4:2-Tc:Mu:Kmr Tn7 (λpir); $Strep^R$. | MB_648 | [29] |
| **Plasmids** | | | |
| pBR-FRT-Kan-FRT2 | pBR322 derivative containing improved FRT-*aph*-FRT cassette, used as template for TransFLP; $Amp^R$, $Kan^R$. | MB_3782 | [18] |
| pBR-FRT-Cat-FRT2 | pBR322 derivative containing improved FRT-*cat*-FRT cassette, used as template for TransFLP; $Amp^R$, $Cm^R$. | MB_3783 | [18] |
| pBR-flp | pBR322 derivative containing FLP+, λ cI857+, λ pR from pCP20 integrated into the *Eco*RV site of pBR322, used for FLP recombination; $Amp^R$ | MB_1203 | [2] |
| pGP704-Sac28 | Suicide plasmid, *oriR6K sacB;* $Amp^R$ | MB_694 | [5] |

| | | | |
|---|---|---|---|
| pGP704-Sac28-[SNP45-T] | pGP704-Sac28 carrying a genome fragment resulting in a site-directed mutation in the SNP45 towards "T" located in the intergenic region between *VCA0106* and *VCA0107* (*vipA*); $Amp^R$ | MB_9062 | This study |
| pGP704-Sac28-[SNP45-G] | pGP704-Sac28 carrying a genome fragment resulting in a site-directed mutation in the SNP45 towards "G" located in the intergenic region between *VCA0106* and *VCA0107* (*vipA*); $Amp^R$ | ND_251 | This study |
| pGP704-Sac28-[SNP45-C] | pGP704-Sac28 carrying a genome fragment resulting in a site-directed mutation in the SNP45 towards "C" located in the intergenic region between *VCA0106* and *VCA0107* (*vipA*); $Amp^R$ | ND_330 | This study |
| pGP704-Sac28-[SNP45-A] | pGP704-Sac28 carrying a genome fragment resulting in a site-directed mutation in the SNP45 towards "A" located in the intergenic region between *VCA0106* and *VCA0107* (*vipA*); $Amp^R$ | ND_331 | This study |
| pGP704-Sac28-SL6Y-[SNP45-G] | pGP704-Sac28 carrying a genome fragment resulting in a site-directed mutation in the SNP45 towards "G" located in the intergenic region between *VCA0106* and *VCA0107* (*vipA*) of strain SL6Y; $Amp^R$ | ND_505 | This study |
| pGP704-Sac28-SP6G-[SNP45-G] | pGP704-Sac28 carrying a genome fragment resulting in a site-directed mutation in the SNP45 towards "G" located in the intergenic region between *VCA0106* and *VCA0107* (*vipA*) of strain SP6G; $Amp^R$ | ND_506 | This study |
| p28-*tfoX* (pGP704-Sac28 -Δ*tfoX*) | pGP704-Sac28 carrying a gene fragment resulting in a deletion within *tfoX* (*VC1153*); $Amp^R$ | MB_1013 | [1] |
| pGP704-28-SacB-Δ*qstR* | pGP704-Sac28 carrying a gene fragment resulting in a deletion within *qstR* (*VC0396*). $Amp^R$. | MB_1118 | [22] |
| pGP704-28-SacB-Δ*tfoY* | pGP704-Sac28 carrying a gene fragment resulting in a deletion within *tfoY* (*VC1722*). $Amp^R$. | MB_4116 | [18] |
| pGP704-Sac28-Δ*vipB* (pGP704-28-SacB-ΔVCA0108) | pGP704-Sac28 carrying a gene fragment resulting in a deletion within *vipB* (*VCA0108*). $Amp^R$. | MB_1123 | [4] |
| pGP704-Sac28-Δ*vasK* (pGP704-28-SacB-ΔVCA0120) | pGP704-Sac28 carrying a gene fragment resulting in a deletion within *vasK* (*VCA0120*). $Amp^R$. | MB_1124 | [4] |
| p28-*hapR* (pGP704-Sac28 -Δ*hapR*) | pGP704-Sac28 carrying a gene fragment resulting in a deletion within *hapR* (*VC0583*). $Amp^R$. | MB_1038 | [1] |
| pGP704-Sac28-ΔIGR276 [SNP45-G] | pGP704-Sac28 carrying a gene fragment resulting in a 276 bp deletion in the intergenic region between *VCA0106* and *VCA0107* (*vipA*) with a SNP45 as "G". 25 bp upstream of *vipA* were kept; $Amp^R$ | ND_559 | This study |
| pGP704-Sac28-ΔIGR276 [SNP45-T] | pGP704-Sac28 carrying a gene fragment resulting in a 276 bp deletion in the intergenic region between *VCA0106* and *VCA0107* (*vipA*) with a SNP45 as "T". 25 bp upstream of *vipA* were kept; $Amp^R$. | ND_560 | This study |
| pGP704-Sac28-ΔIGR336 [SNP45-G] | pGP704-Sac28 carrying a gene fragment resulting in a 336 bp deletion in the intergenic region between *VCA0106* and *VCA0107* (*vipA*) with a SNP45 as "G". 25 bp upstream of *vipA* were kept; $Amp^R$ | ND_561 | This study |
| pGP704-Sac28-ΔIGR336 [SNP45-T] | pGP704-Sac28 carrying a gene fragment resulting in a 336 bp deletion in the intergenic region between *VCA0106* and *VCA0107* (*vipA*) with a SNP45 as "T". 25 bp upstream of *vipA* were kept; $Amp^R$ | ND_562 | This study |

| | | | |
|---|---|---|---|
| pGP704-Sac28-ΔIGR396 (Δfull) | pGP704-Sac28 carrying a gene fragment resulting in a 396 bp deletion of the intergenic region between *VCA0106* and *VCA0107* (*vipA*) – including SNP45. 10 bp downstream of *VCA0106* and 25 bp upstream of *vipA* were kept; $Amp^R$ | ND_368 | This study |
| pGP704-Sac-Kan | Suicide plasmid, *oriR6K sacB*, $Kan^R$. | MB_6038 | [6] |
| pGP704-Sac-Kan-L6G- [SNP45-G] | pGP704-Sac-Kan carrying a genome fragment resulting in a site-directed mutation in the SNP45 towards "G" located in the intergenic region between *VCA0106* and *VCA0107* (*vipA*) of strain L6G; $Kan^R$ | ND_503 | This study |
| pGP704-Sac-Kan-SL5Y- [SNP45-G] | pGP704-Sac-Kan carrying a genome fragment resulting in a site-directed mutation in the SNP45 "G" located in the intergenic region between *VCA0106* and *VCA0107* (*vipA*) of strain SL5Y; $Kan^R$ | ND_506 | This study |
| pGP704-Sac-Kan-SL4G- [SNP45-G] | pGP704-Sac-Kan carrying a genome fragment resulting in a site-directed mutation in the SNP45 towards "G" located in the intergenic region between *VCA0106* and *VCA0107* (*vipA*) of strain SL4G; $Kan^R$ | ND_507 | This study |
| pGP704-Sac-Kan-ΔIG60bp [SNP45-T] | pGP704-Sac-Kan carrying a genome fragment resulting in a ~60bp deletion in the intergenic region between *VCA0106* and *VCA0107* (*vipA*) with a SNP45 as "T". 10bp downstream of *VCA0106* and 10bp upstream of *vipA* were kept; $Kan^R$ | ND_754 | This study |
| pGP704-Sac-Kan-ΔIG60 [SNP45-G] | pGP704-Sac-Kan carrying a genome fragment resulting in a ~60bp deletion in the intergenic region between *VCA0106* and *VCA0107* (*vipA*) with a SNP45 as "G". 10bp downstream of *VCA0106* and 10bp upstream of *vipA* were kept; $Kan^R$ | ND_763 | This study |
| pGP704-Sac-Kan-chg[-10box] [SNP45-T] | pGP704-Sac-Kan carrying a genome fragment resulting in a site-directed mutation in the -10 element (AA to GC) located in the intergenic region between *VCA0106* and *VCA0107* (*vipA*) with a SNP45 as "T"; $Kan^R$ | ND_756 | This study |
| pGP704-Sac-Kan-chg[-10box] [SNP45-G] | pGP704-Sac-Kan carrying a genome fragment resulting in a site-directed mutation in the -10 element (AA to GC) located in the intergenic region between *VCA0106* and *VCA0107* (*vipA*) with a SNP45 as "G"; $Kan^R$ | ND_767 | This study |

*reference locus tags belong to reference strain N16961 according to [30].

**Table S4.** Statistics on PacBio whole-genome sequencing and genome assembly of strain ATCC25872.

| | **ATCC25872** |
|---|---|
| **Strain ID** | MB#276 |
| **BioSample ID** | SAMN13736322 |
| **GenBank accession numbers (chr 1 / chr 2)** | CP047305 / CP047306 |
| **Number of bases** | 7.73 Gbp |
| **Number of reads** | 414,427 |
| **Mean read length** | 18,897 bp |
| **Total number of contigs** | 2 (chr1+chr2) |
| **Contig length after circularization** | 2,945,491 bp (chr1)<br>1,072,958 bp (chr2) |
| **Total genome size** | 4,018,449 bp |
| **Mean coverage** | 1,796 x |
| **GC content** | 47.8% (chr1)<br>46.9% (chr2) |
